# Ablation of ZC3H11A causes early embryonic lethality and dysregulation of metabolic processes

**DOI:** 10.1101/2022.09.14.508037

**Authors:** Shady Younis, Alice Jouneau, Mårten Larsson, Jean-Francois Oudin, Vincent Brochard, Leif Andersson

## Abstract

ZC3H11A is a stress-induced mRNA binding protein required for efficient growth of nuclear-replicating viruses, while being dispensable for the viability of cultured human cells. The cellular functions of ZC3H11A during embryo development are unknown. Here we report the generation and phenotypic characterization of *Zc3h11a* knock-out mice. Heterozygous null *Zc3h11a* mice were born at the expected frequency without distinguishable phenotypic differences compared with wild-type. In contrast, homozygous null *Zc3h11a* mice were missing, indicating that *Zc3h11a* is crucial for embryonic viability and survival. *Zc3h11a*^−/–^ embryos were detected at the expected Mendelian ratios up to late preimplantation stage (E4.5). However, phenotypic characterization at E6.5 revealed degeneration of *Zc3h11a*^−/–^ embryos, indicating developmental defects around the time of implantation. Transcriptomic analyses documented a dysregulation of glycolysis and fatty acid metabolic pathways in *Zc3h11a*^−/–^ embryos at E4.5. Proteomic analysis indicated a tight interaction between ZC3H11A and mRNA-export proteins in embryonic stem cells. Furthermore, CLIP-seq analysis demonstrated that ZC3H11A binds a subset of mRNA transcripts that are critical for metabolic regulation of embryonic cells. Altogether, the results show that ZC3H11A is participating in export and post-transcriptional regulation of selected mRNA transcripts required to maintain metabolic processes in embryonic cells. While ZC3H11A is essential for the viability of the early mouse embryo, inactivation of *Zc3h11a* expression in adult tissues using a conditional knock-out did not lead to obvious phenotypic defects.

## Introduction

The zinc finger CCCH domain-containing protein 11A (ZC3H11A) is a stress-induced mRNA-binding protein that is required for the efficient growth of several human nuclear replicating viruses, including human immunodeficiency virus (HIV-1), influenza A virus (IAV), human adenovirus (HAdV) and herpes simplex virus 1 (HSV-1) [1]. Proteomic studies on human cells have indicated that ZC3H11A is a component of the transcription-export (TREX) complex [2]. Functional studies indicated that ZC3H11A selectively export newly transcribed viral mRNAs to the cytoplasm during virus infection [1, 3]. Thereby, inactivation of ZC3H11A in human cells impaired the export of a subset of viral mRNA transcripts and resulted in a dramatic reduction in virus growth [1]. These important functions of ZC3H11A in the growth cycle of several human viruses makes ZC3H11A a potential target for development of an anti-viral therapy. The aim of the present study was to develop an animal model to study the molecular functions of ZC3H11A in prenatal and postnatal development.

The TREX complex serves a key function in nuclear mRNA export and consists of multiple conserved core subunits including ALYREF (RNA binding adaptor of TREX), UAP56 (DEAD-box type RNA helicase) and a stable subcomplex called THO, which in turn consists of at least six subunits [4, 5]. Proteomic studies using human cells have indicated that ZC3H11A is an auxiliary component of the TREX complex, but did not consider it as a core subunit of the TREX complex [6, 7]. THO proteins are conserved from yeast to human and play pivotal roles during embryo development, cell differentiation and cellular response to stimuli [8, 9]. It has been reported that the disruption of THO proteins, such as THOC1, THOC2 or THOC5, leads to early embryonic lethality [9–11]. The TREX complex controls the mRNA export in a selective manner, where individual TREX components appear to be required for export of distinct subsets of mRNAs [12]. For instance, THOC2 or THOC5 are required for the export of mRNAs essential for pluripotency such as *Nanog, Sox2* and *Klf4* in mouse embryonic stem cells [9]. Despite several reports characterizing the role of THO proteins during embryogenesis, the cellular function of ZC3H11A during embryo development is unknown.

In the current study, we established *Zc3h11a* knock-out (KO) mouse models to study the effect of *Zc3h11a* loss of function on embryo development. Our results identify ZC3H11A as a fundamental protein required for early embryo growth. Disruption of ZC3H11A is homozygous lethal and leads to complete failure of embryo development and survival. Using proteomic and RNA-seq analyses, we show that the ZC3H11A protein interacts with TREX complex core proteins in mouse embryonic stem cells. ZC3H11A is apparently an auxiliary factor participating in export and post-transcription coordination of selected mRNA transcripts required to maintain the metabolic processes in embryonic cells. Interestingly, *Zc3h11a* inactivation in adult mouse tissues using an inducible mouse model showed that the ZC3H11A protein is dispensable for postnatal tissue growth.

## Results

### *Zc3h11a* inactivation in mice is lethal in the homozygous condition

*Zc3h11a* is located on chromosome 1 in both human and mouse genomes and harbors the coding sequence of another gene encoding the DNA-binding zinc-finger protein ZBED6 [13–18] (Figure 1A). We used two strategies to target the *Zc3h11a* coding exons without affecting *Zbed6*. The first *Zc3h11a* mouse model was developed by targeting exon 3 using the CRISPR/cas9 system with two guide RNAs flanking the targeted sequences. This resulted in both a deletion of 567 bp including the entire exon 3 and a frameshift (Figure 1B). The second mouse model was developed by inserting loxP sites flanking the coding sequence of exon 2 using homologous recombination (Figure 1C). These loxP mice were crossed with mice expressing Cre recombinase in germ-line (PGK-Cre), which resulted in a deletion of 1.5 kb containing exon 2 and removal of the zinc finger domains of the encoded ZC3H11A protein (Figure 1C). For each model, heterozygous mice were crossed and the offspring were genotyped. No *Zc3h11a*^*–/–*^ mice were obtained from heterozygous matings (Figure 1D and E), with the exception of one single homozygous *Zc3h11a*^*–/–*^ female from the loxP mouse model (1 out of 204 mice). When we crossed this KO female with *Zc3h11a*^*+/–*^ males, 10 out of 10 progeny were heterozygous *Zc3h11a*^*+/–*^. The probability to get this outcome if both parents are heterozygous is P=0.5^10^=0.001. The result confirms our interpretation that one single homozygous KO survived and were fertile.

**Figure 1.**
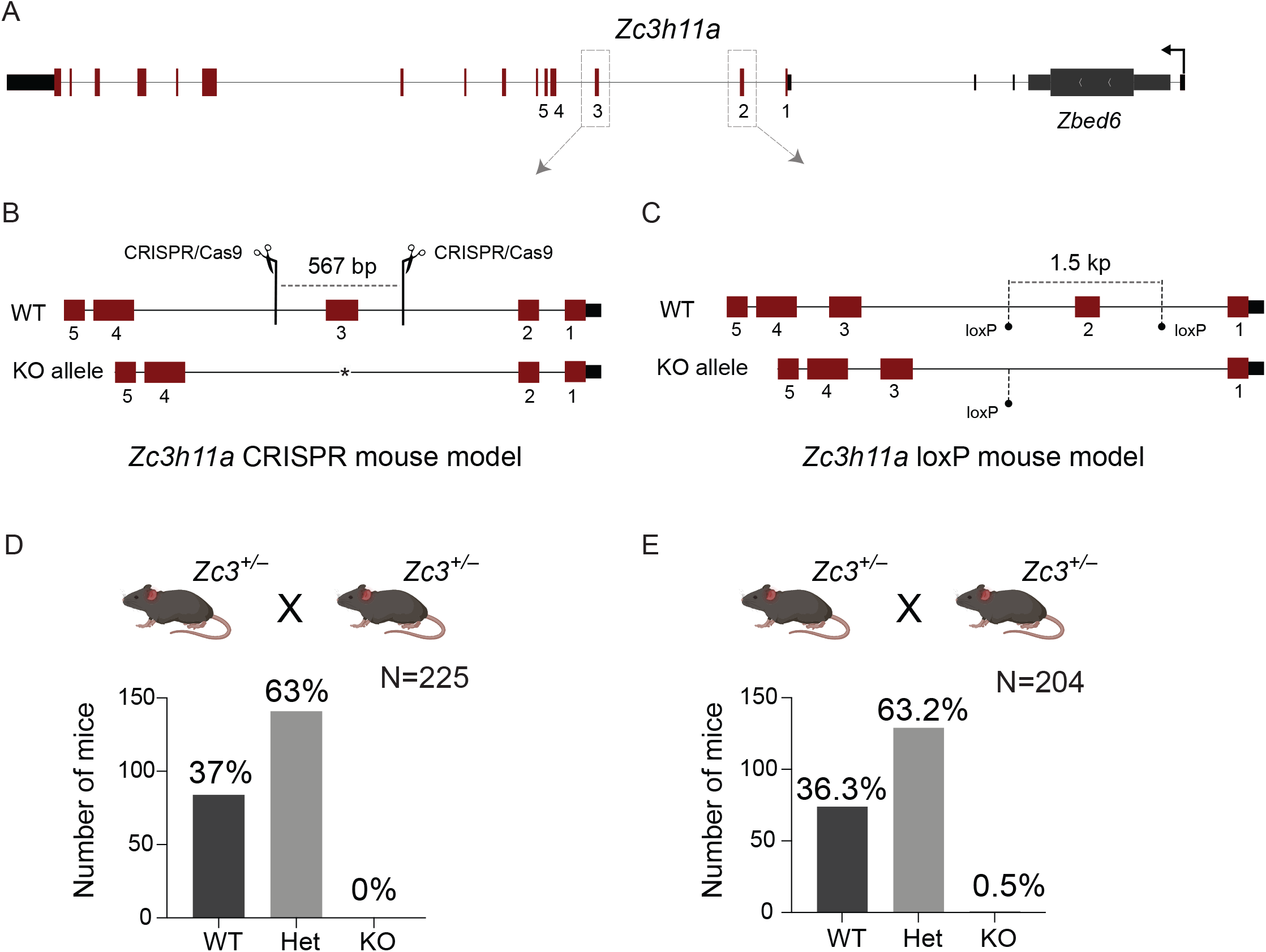
Development of *Zc3h11a*^−/–^ mouse models. (A) The *Zc3h11a* locus showing the targeted exons for generating *Zc3h11a*^−/–^ mouse models. (B) Two CRISPR/cas9 guide RNAs were used to delete exon 3 of *Zc3h11a*. Scissors indicate the location of the gRNAs and the length of deleted sequences. (C) Two homology arms were used to insert loxP sites flanking exon 2. The conditional knock-out mice were crossed with mice expressing Cre in germ line to eliminate the sequences between the loxP sites resulting in the elimination of the entire exon 2 coding sequences. (D and E) Genotyping of the offspring of *Zc3h11a* heterozygous matings (*Zc3*^+/–^ X *Zc3*^+/–^) using the CRISPR/cas9-based KO mouse model (D) and the loxP/Cre-based KO mouse model (E). The total numbers of genotyped mice at week 4 are indicated.

### *Zc3h11a* deletion results in embryonic degeneration

In order to explore at what point ZC3H11A is essential for embryo survival, we collected and genotyped embryos at different time points post *Zc3h11a*^*+/–*^ X *Zc3h11a*^*+/–*^ mating (Figure 2A). The genotyping of embryos at embryonic day E4.5 prior to implantation revealed expected Mendelian proportions (Figure 2A). However, a clear deviation from expected Mendelian proportions was observed after implantation (Figure 2A, bottom panel). Remarkably, phenotyping at E6.5 showed dramatic changes in the morphology of the *Zc3h11a*^*–/–*^ embryos with a large degree of tissue degeneration, whereas *Zc3h11a*^*+/–*^ heterozygotes appeared morphologically indistinguishable from the WT embryos (Figure 2B).

**Figure 2.**
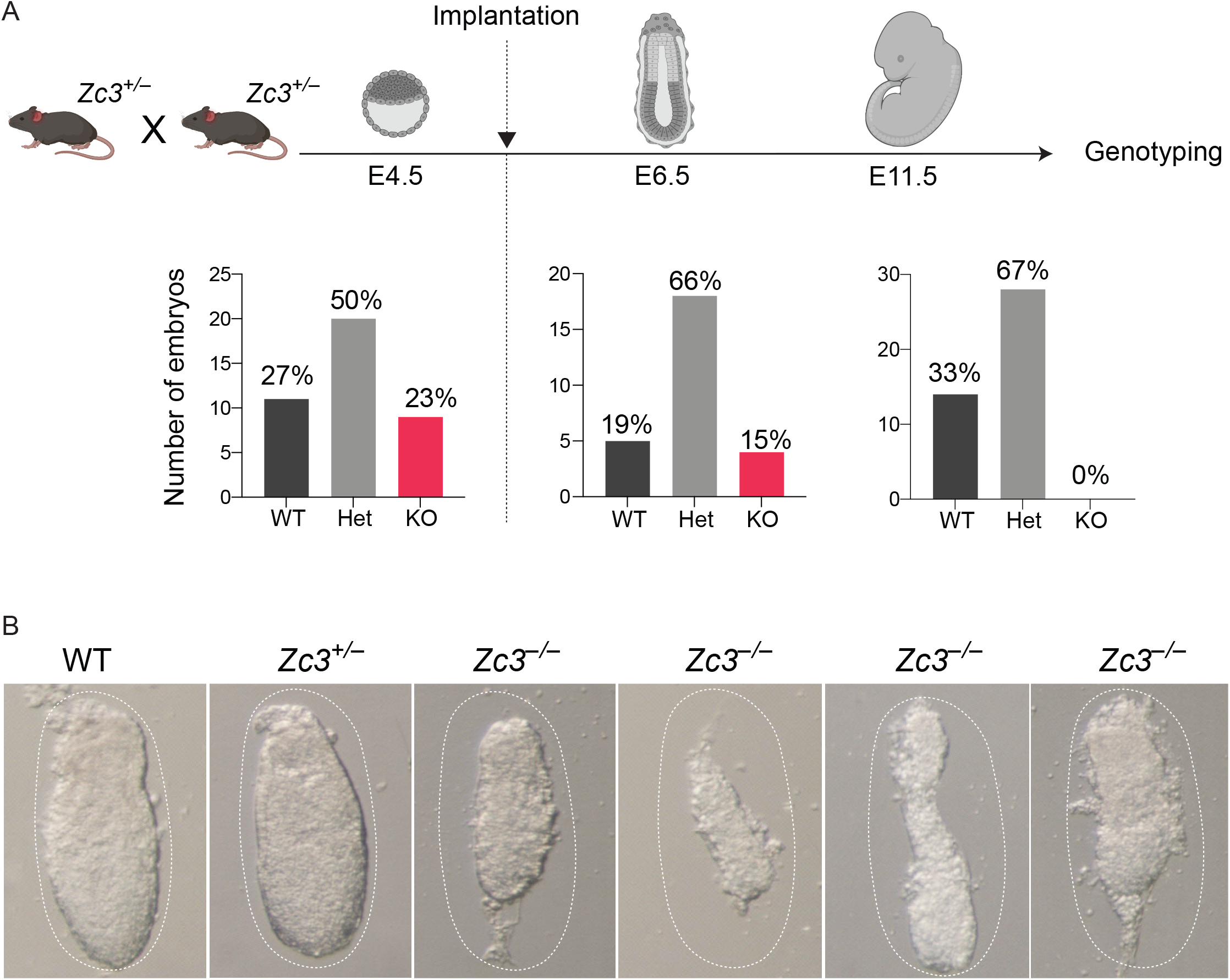
Ablation of *Zc3h11a* leads to early embryo degeneration. (A, top) Schematic illustration showing embryo stages and time points of collecting embryos for genotyping of *Zc3h11a*. (A, low) PCR genotyping of collected embryos at the above time points. (B) Morphology of collected embryos at E6.5 from *Zc3h11a* heterozygous mating (*Zc3*^+/–^ X *Zc3*^+/–^).

### ZC3H11A is highly expressed at early stages of embryonic development

The lethal effect of *Zc3h11a* inactivation in mouse embryos encouraged us to explore the cellular localization of ZC3H11A at early embryonic stages. We used immunofluorescence (IF) staining to visualize the ZC3H11A protein and the nuclear speckles marker SRSF2 (SC35) for expression profiling in mouse 2-cell and blastocyst stages. The IF analysis indicated that ZC3H11A was expressed at a detectable level as early as the 2-cell stage, with clear nuclear localization (Figure 3A, top panel). The z-stack imaging of the blastocysts showed that ZC3H11A was expressed in trophectoderm (Figure 3A, middle panel) as well as in inner cell mass (ICM) (Figure 3A, bottom panel). The localization pattern of ZC3H11A in ICM was overlapping with the nuclear speckles as indicated using the anti-SC35 antibody (Figure 3A and B). This subcellular localization of ZC3H11A in mouse embryonic cells is similar to the ZC3H11A localization in human cell lines [1]. Re-analyzing single cell RNA-seq data from Deng *et. al*. [19] revealed that *Zc3h11a* mRNA is highly expressed in mouse embryos as early as the zygotic stage, indicating maternal contribution (Figure 3C).

**Figure 3.**
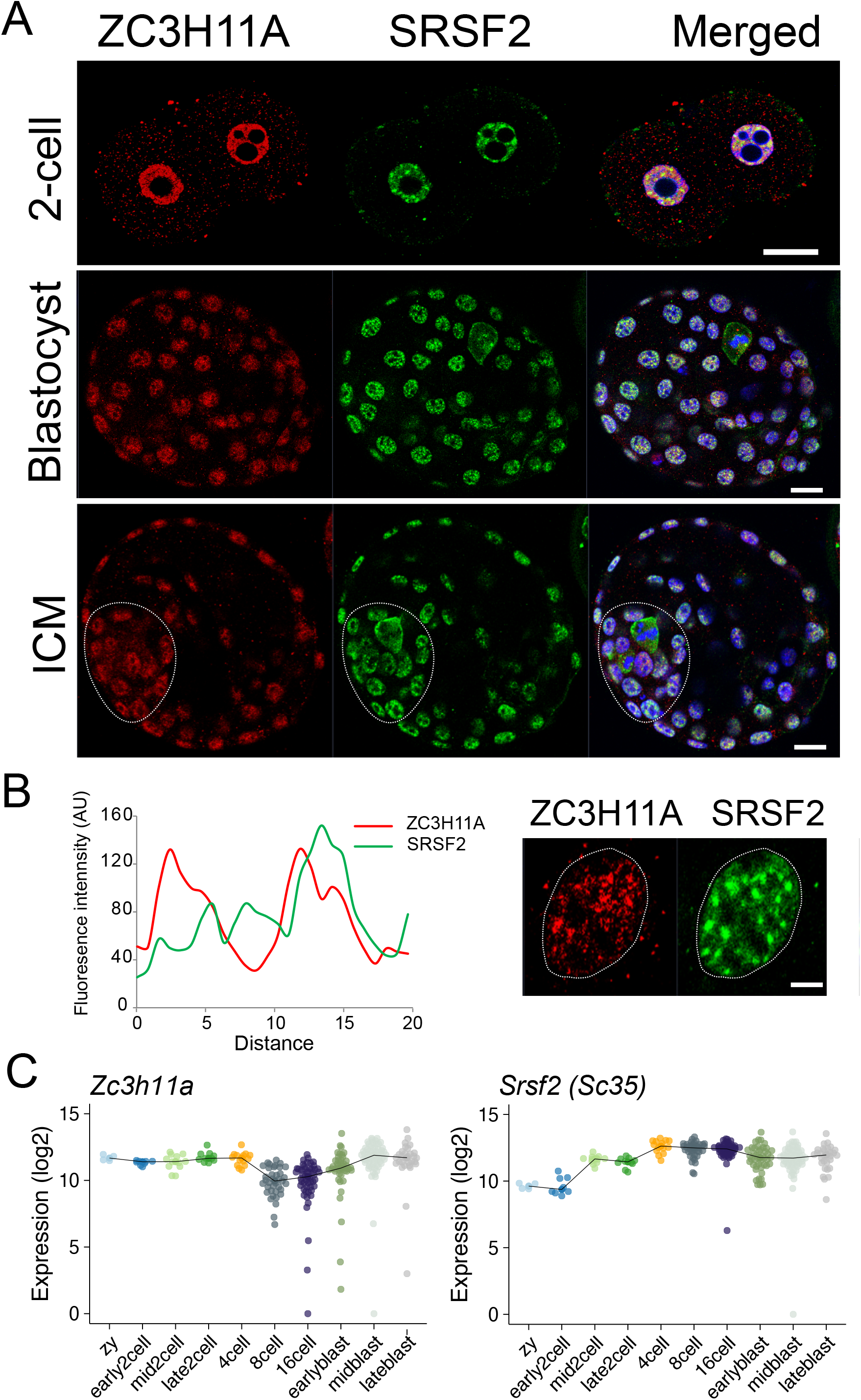
Cellular localization of ZC3H11A in early embryonic cells. (A) Immunofluorescence staining of mouse embryos using anti-ZC3H11A and anti-SRSF2 (SC35) antibodies at 2-cell (top) and blastocyst stage (middle and bottom). Middle panel shows a z-projection of the whole blastocyst while the bottom panel is a mid-section through the ICM. (Scale bar: 20 μm). (B) Fluorescent intensity profile of the ZC3H11A signal and paraspeckle marker SRSF2 signal across paraspeckles in an ICM nucleus showing the co-localization of ZC3H11 to paraspeckles. (Scale bar: 5 μm.). (C) Expression of *Zc3h11a* and *Srsf2* measured by smartseq2 single cell RNA-seq. Re-analysis of data from Deng *et. al*. [19].

### Disrupted metabolic pathways in the *Zc3h11a*^*–/–*^ embryos

The degeneration of *Zc3h11a* ^*–/–*^ embryos during early embryo development (E6.5) encouraged us to perform whole transcriptome analysis of stage E4.5 embryos to reveal the dysregulated pathways that led to the degeneration of *Zc3h11a* ^*–/–*^ embryos at E6.5. We collected embryos from *Zc3h11a*^*+/–*^ X *Zc3h11a*^*+/–*^ matings and extracted the RNA from the embryonic part for sequencing (Figure 4A, left). Principle component analysis (PCA) of RNA-seq data showed that Het (*Zc3h11a*^*+/–*^) and WT (*Zc3h11a*^*+/+*^) embryos clustered together and apart from the KO (*Zc3h11a*^*–/–*^) embryos (Figure 4A). This result is in agreement with the observed morphological similarity between WT and Het embryos (Figure 2B). Furthermore, the differential expression (DE) analysis between WT and Het did not detect any significant DE genes with FDR <0.05. Therefore, we performed the DE analysis between KO embryos vs. WT and Het embryos that revealed 660 DE genes (FDR <0.05) out of ~11,000 expressed genes (Table S1). Among these DE genes, 419 were up-regulated and 241 were down-regulated in KO embryos (FDR <0.05). Next, we performed a gene set enrichment analysis (GSEA) using the DE genes in KO embryos in order to further explore the function of ZC3H11A. The GSEA of ranked DE genes in KO embryos using the hallmark gene sets revealed a significant negative enrichment (FDR <0.05) of genes involved in glycolysis, fatty acid metabolism pathways and epithelial-mesenchymal transition (EMT) processes (Figures 4B-4D). The heatmaps present the expression of the subset of genes that contributed the most to the indicated pathway enrichment among significantly down-regulated genes in KO embryos (Figures 4B-4D). Among the key down-regulated genes, contributing to the significant GSEA result, are lactate dehydrogenase A (*Ldha*), which has an essential role in glycolysis, and disruption of *Ldha* causes congenital disorders of carbohydrate metabolism [20, 21]; enoyl-CoA hydratase and 3-hydroxyacyl CoA dehydrogenase (*Ehhadh*), which is involved in fatty acid beta-oxidation using acyl-CoA oxidase [22, 23]; and dickkopf WNT signaling pathway inhibitor 1 (*Dkk1*), which is involved in several processes including cell fate determination and cell differentiation processes during embryogenesis [24]. On the other hand, the positively enriched gene sets among up-regulated genes in the KO mice included genes in the P53 pathway and autophagy process-related genes (Figures 5A, 5B and S1). This includes the up-regulation of autophagy related 12 (*Atg12*) and microtubule-associated proteins 1A/1B light chain 3A (*Map1lc3a*) genes (Figure 5E). MAP1LC3A is known as LC3A protein and is required for autophagosome formation [25].

**Figure 4.**
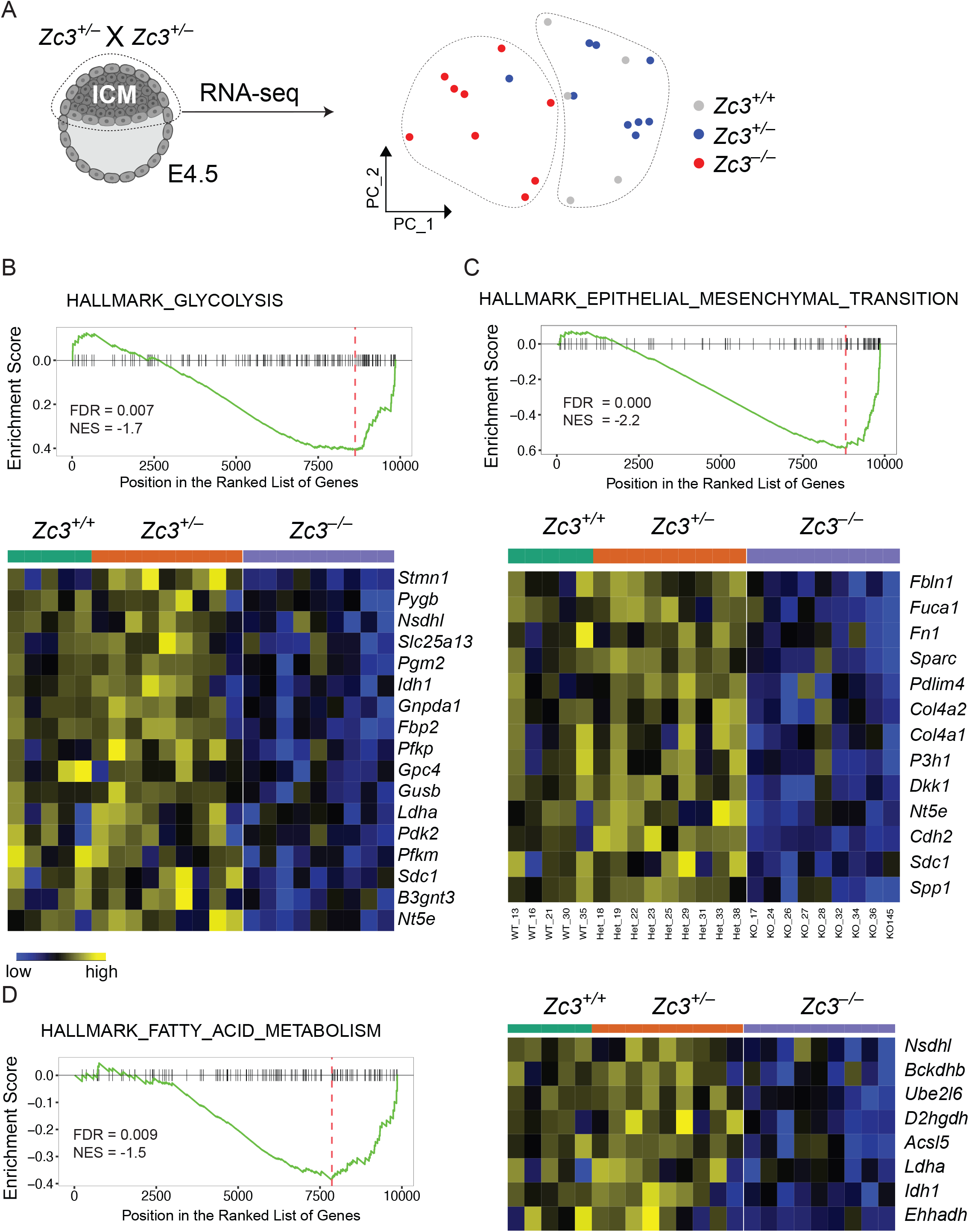
Transcriptome analysis reveals dysregulated pathways in *Zc3h11a*^−/–^ embryos. (A, left) Dissected inner cell mass (ICM) cells were used for RNA-sequencing. (A, right) Principle component analysis (PCA) of RNAseq data from embryonic parts at E4.5. Dots represent individual embryos and colors represent different genotypes. (B-D) Gene set enrichment analysis (GSEA) of ranked DE genes in *Zc3h11a*^−/–^ embryos using hallmark gene sets. (B-C, below) Heatmaps showing the expression of the genes contributing to the above pathways and found significantly down-regulated in *Zc3h11a*^−/–^ embryos (FDR <0.05). (E) Heatmap showing the expression of the genes contributing to fatty acid metabolism pathway and significantly down-regulated in *Zc3h11a*^−/–^ embryos (FDR <0.05). FDR: false discovery rate, NES: normalized enrichment score.

**Figure 5.**
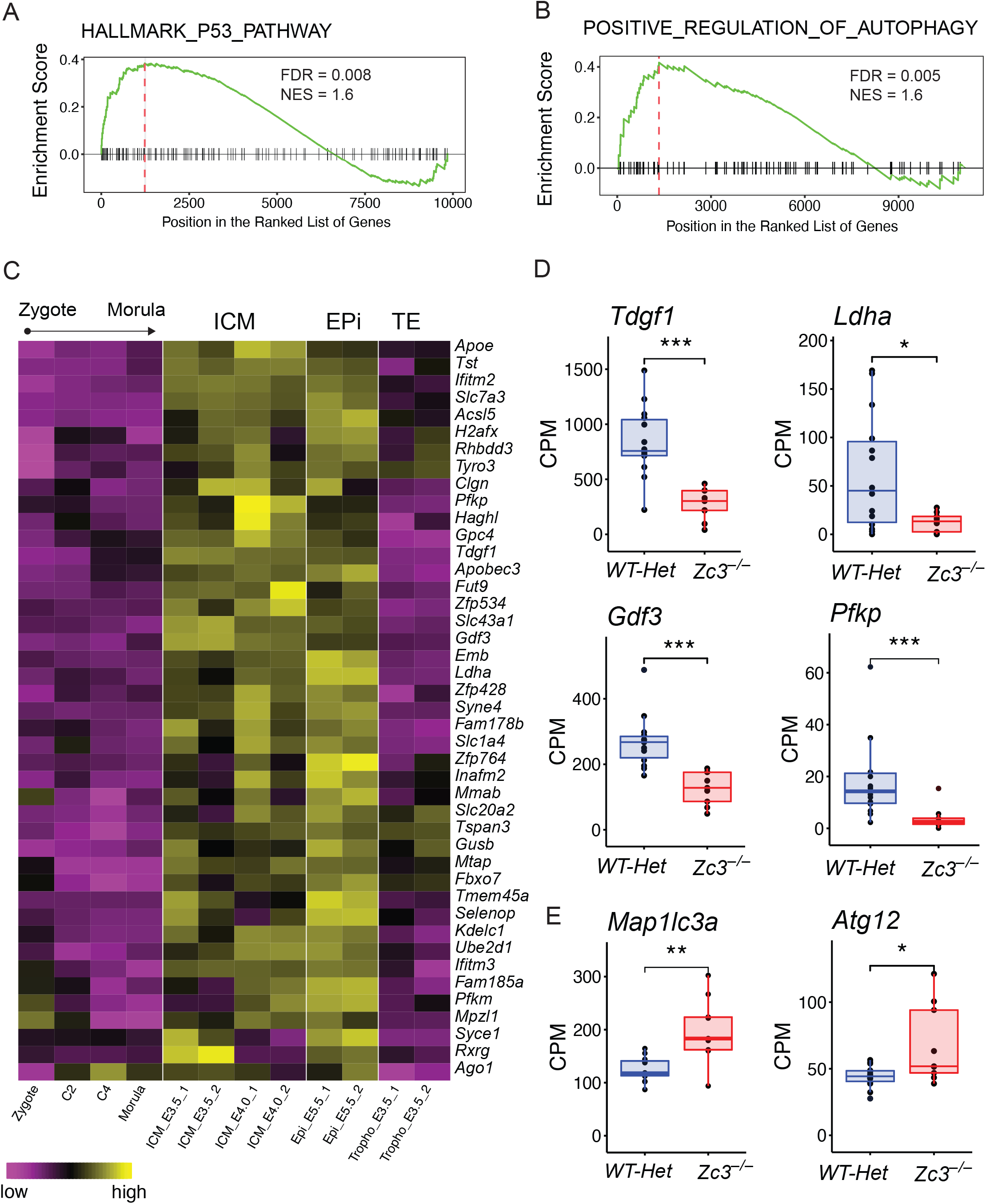
Down-regulated genes in *Zc3h11a*^−/–^ embryos are ICM-related. (A-B) GSEA of ranked DE genes in *Zc3h11a*^−/–^ embryos with positive enrichment for the P53 pathway (A) and autophagy-related genes (B) among the up-regulated genes in *Zc3h11a*^−/–^ embryos (FDR <0.05). Heatmap of down-regulated genes in *Zc3h11a*^−/–^ embryos (FDR <0.05) and their expression profile during embryonic stages as indicated. Re-analyzed data from GSE76505 and E-MTAB-2950. (D) Expression level of the indicated genes as count per millions (CPM). *, ** and *** correspond to *FDR*< 0.05, 0.01 and 0.001, respectively.

To get further insight on which cell type in blastocysts was most affected by *Zc3h11a* inactivation, we explored the expression profile of the DE genes in *Zc3h11a*^−/–^ embryos in embryonic lineages (ICM and epiblast) and trophectoderm (TE). Using previously published datasets of mouse gene expression (GSE76505 [26]), the ICM/TE ratio of expression was computed for genes down-regulated and up-regulated in the KO embryos (FDR ≤0.05, fold change ≥2). We also explored the expression of DE genes at earlier stages using gene expression dataset (E-MTAB-2950) [27]. This showed that down-regulated genes in *Zc3h11a*^−/–^ embryos are primarily expressed in the ICM/early epiblast rather than trophectoderm (Figure 5C), while the expression of the up-regulated genes is nearly equally present in ICM/early epiblast and TE (Figure S2). The down-regulated genes in *Zc3h11a*^−/–^ embryos with high expression in the ICM include: *Ldha*, teratocarcinoma-derived growth factor (*Tdgf1, Cripto)*, growth differentiation factor 3 (*Gfd3)*, phosphofructokinase (*Pfkp*) (Figure 5D). GDF3 is an analog of NODAL and uses TDGF1 as co-factor [28]. Both *Ldha, Pfkp, Pfkm* and *Pdk2* are down-regulated in *Zc3h11a*^−/–^ embryos and are involved in glycolysis and lactate production, as indicated in the GSEA (Figure 4). At peri-implantation stages, there is a major metabolic switch from oxidative phosphorylation to anaerobic glycolysis, with increased lactate production [29, 30]. *Cripto*/*Tdgf1* has been reported as an essential factor regulating this metabolic switch [31]. That explains the GSEA results that show that the down-regulated pathways mostly concern metabolic regulation processes. Altogether, this strongly suggests that the primary consequence of ZC3H11A deficiency is in the ICM, due to perturbed metabolic regulation. The enrichment of genes associated with autophagy and apoptosis-related pathways (Figure 5A-5B) among the up-regulated genes in *Zc3h11a*^*–/–*^ embryos could be secondary effect caused by the metabolic stress encountered by the ICM cells [32, 33].

### ZC3H11A is associated with the RNA export machinery in embryonic stem cells

In human somatic cells, ZC3H11A has been recently characterized as an RNA-export protein that functions through its interaction with TREX complex proteins [1]. In order to identify its interacting partners in embryonic cells and to investigate if ZC3H11A maintains its association with the TREX complex in mouse embryonic stem cells (mESCs), we performed co-immunoprecipitations (co-IPs) using anti-ZC3H11A, anti-THOC2 and anti-IgG antibodies followed by mass spectrometry (MS) analyses (Figure 6A). Statistical analyses of detected MS intensities from the biological replicates (n=4) revealed a number of proteins with statistically significant interaction with ZC3H11A and THOC2 (Figure 6B). Proteins belonging to the TREX complex and RNA export machinery are highlighted in bold. The log-fold change in protein intensities in the ZC3H11A co-IP relative to the IgG co-IP is presented along with the adjusted *P*-values (Figure 6C). The interaction between ZC3H11A and THOC2 was validated by a reciprocal co-IP and western blot using mESCs (Figure 6D). The majority of the significant partners interacting with ZC3H11A are part of the TREX complex and also showed significant enrichment in the THOC2 co-IP, including THOC5, THOC7 (Figure 6E), THOC1 and THOC6 (Figure 6C). ZC3H11A also interacts with other RNA-binding proteins that are required for RNA maturation, such as polyadenylate-binding nuclear protein 1 (PABPN1) [34]; FYTTD1, which acts as an adaptor for RNA helicase UAP56 [35]; and the RNA export adaptor ALYREF/THOC4 [36] (Figure 6D and F). Notably, almost half of the ZC3H11A partners detected by co-IP were also found in the THOC2 co-IP (Figure S3). These data indicate that ZC3H11A is an essential component of the TREX complex that is known to play pivotal roles during embryogenesis and for maintaining pluripotency of ESCs [9, 10]. Furthermore, the proteomic analysis identified additional interacting partners, independent of the TREX complex, such as the RNA-binding protein DDX18; the PRC2 components SUZ12 and JARID2; and the two zinc finger proteins ZNF638 and ZFP57 (Figure 6G and S3B). DDX18 is an RNA binding protein that plays a crucial role in pluripotency and self-renewal of embryonic stem cells [37].

**Figure 6.**
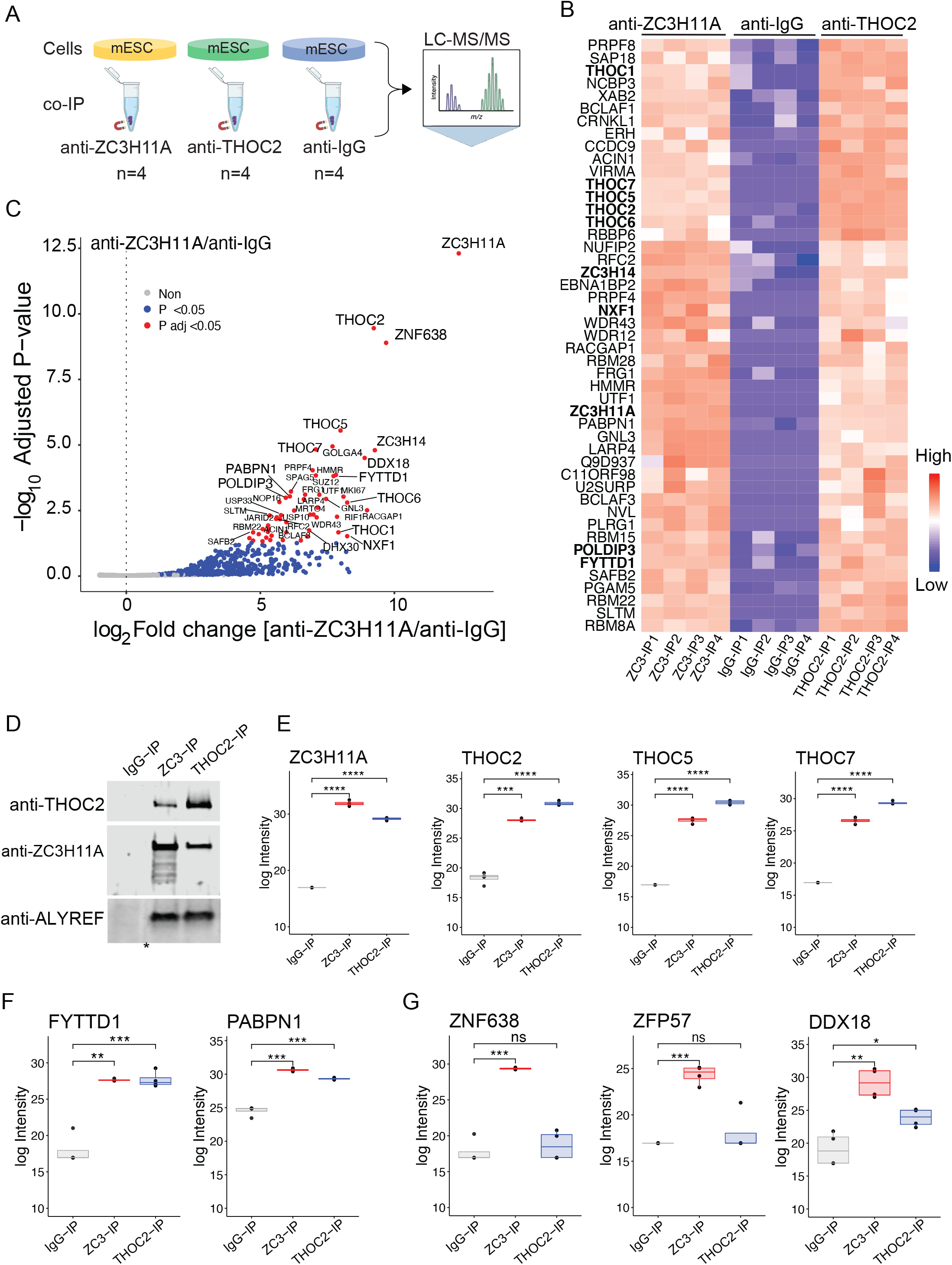
ZC3H11A binds RNA-export TREX complex proteins in mESC. (A) Schematic illustration of co-immunoprecipitation (co-IP) mass-spectrometry experiments using anti-ZC3H11A, anti-THOC2 and anti-IgG antibodies and mouse embryonic stem cells (mESCs). (B) Heatmap of the interacting partners to ZC3H11A (adjusted *P* <0.05). Data presented as log intensities of four replicates. Proteins associated with the TREX complex and mRNA export are in bold. (C) Volcano plot showing the enrichment of co-IP proteins from anti-ZC3H11A/anti-IgG. Western blot of reciprocal co-IP using anti-ZC3H11A, anti-THOC2 and anti-IgG antibodies and probed with the indicated antibodies. Asterisk indicates a cut in the western blot membrane. Log intensities of the ZC3H11A and THOC proteins. (F) Log intensities of FYTTD1 (UAP56) and the polyadenylation factor PABPN1. (G) Log intensities of proteins interacting with ZC3H11A independent of THOC2 and the TREX complex. *, **,*** and **** correspond to adjusted *P* <0.05, 0.01, 0.001 and 0.0001, respectively. ns: not significant.

### ZC3H11A selectively binds mRNA transcripts in mESCs

Previous studies using human somatic cells indicated that ZC3H11A is an RNA-binding protein that selectively binds subsets of mRNA upon stress or viral infection [1]. To study the RNA-binding properties of ZC3H11A in embryonic cells, we performed UV-crosslinking of mESCs followed by ZC3H11A immunoprecipitation (CLIP) and RNaseI treatment to isolate the RNA protected by ZC3H11A. We used two anti-ZC3H11A antibodies to minimize any artifact caused by antibodies, and anti-ALYREF and anti-IgG as positive and negative controls, respectively. High-throughput sequencing of the RNA isolated by CLIP (CLIP-seq) revealed an almost exclusive interaction between ZC3H11A and protein-coding mRNAs in mESCs (Figure S4A), with a preference to bind 3’UTRs over the 5’UTRs (Figure 7A). The analysis of ZC3H11A CLIP-seq peaks from the two ZC3H11A antibodies revealed a significant enrichment of short purine-rich motifs (Figure 7B, top panel). Moreover, ZC3H11A exhibited strong binding to the paraspeckle *Neat1* transcript (Figure S4B), similar to what has been observed in human somatic cells [1]. Comparing the CLIP-seq ZC3H11A mRNA targets with genes that were significantly down-regulated in RNA-seq data, we identified subsets of genes as putative direct targets of ZC3H11A in mESCs (Figure 7B, bottom panel). The gene ontology analysis of these genes suggested that they are involved in germ cell development and metabolic processes (Figure 7C). These 29 genes were dramatically down-regulated in *Zc3h11a*^*–/–*^ embryos (Figure 7D) and are involved in cellular processes vital for embryonic development [38–41]. Putative direct targets included the *Tdgf1*, nucleoporin 85 *(Nup85)*, proliferation-associated protein 2G4 *(Pa2g4)* and gap junction protein beta 3 *(Gjb3)* genes. The CLIP-seq analysis detected ZC3H11A binding sites at the 3’UTR of these genes that were either overlapped with ALYREF binding sites (for *Tdgf1 and Pa2g4*) or distinct from them in *Nup85* (Figure 7E). These results suggest a crucial role of ZC3H11A in post-transcriptional processing and mRNA export of key genes in embryonic cells.

**Figure 7.**
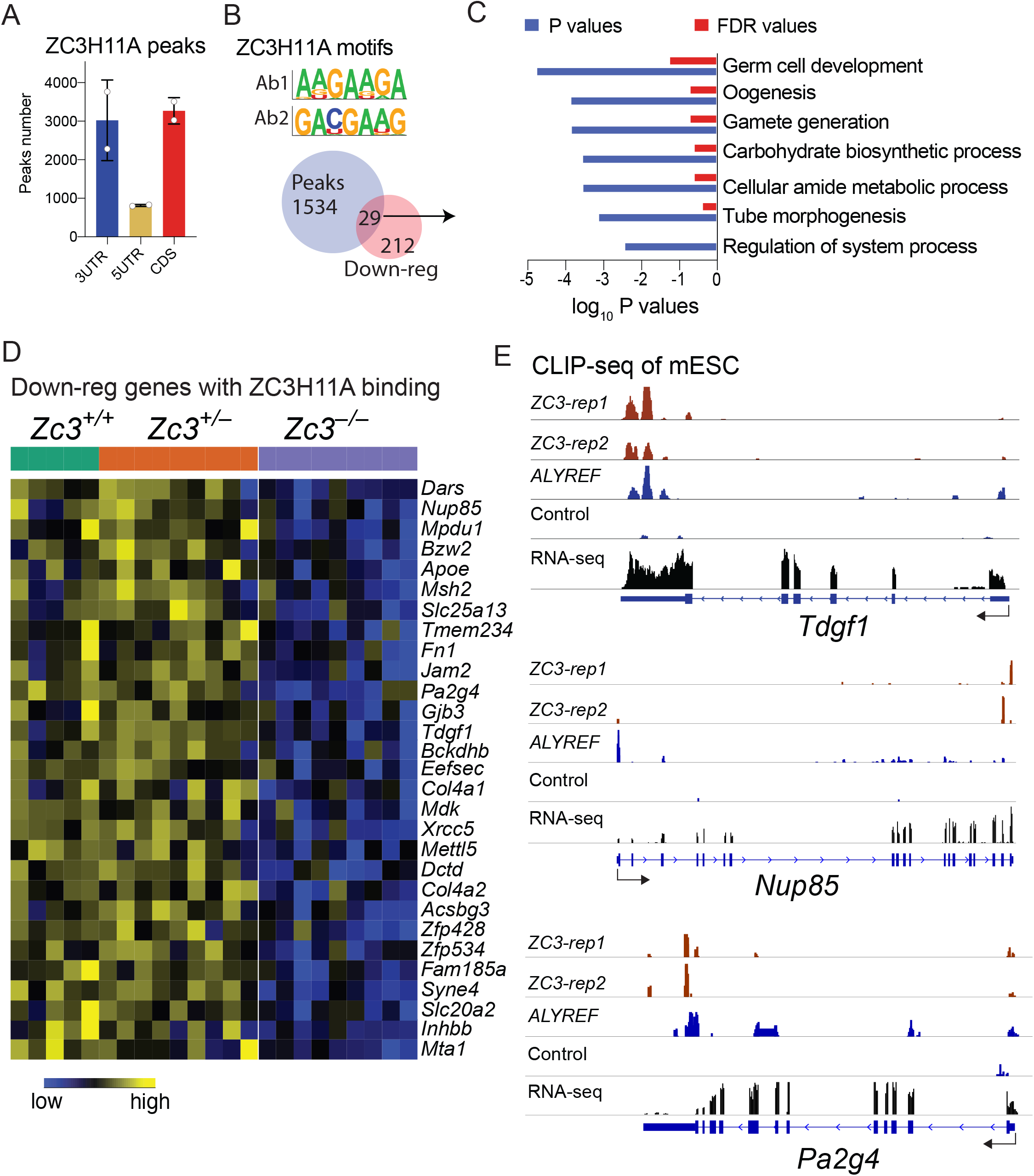
CLIP-seq analysis of ZC3H11A RNA targets in mESCs. (A) Distribution of the proportion of ZC3H11A CLIP-seq mapped reads over the various elements of a gene in mESC using two anti-ZC3H11A antibodies and an anti-IgG control antibody. (B, top) Predicted motifs for ZC3H11A binding. (B, bottom) The overlap between differential down-regulated genes in KO embryos and predicted ZC3H11A CLIP-seq targets. (C) Gene ontology analysis of the down-regulated genes with ZC3H11A binding sites. (D) Heatmap of the down-regulated genes with ZC3H11A binding sites. (E) The visualization of CLIP-seq reads and their distribution over the indicated genes. Black arrows indicate the direction of transcription from 5’ UTR to 3’ UTR.

### ZC3H11A is required for *in vitro* derivation of ESCs

To further understand the role of ZC3H11A in the peri-implantation development and especially its role in the pluripotent epiblast, twenty-five E3.5 blastocysts were recovered from matings between heterozygous mice, and cultured *in vitro*. From these, 14 ESC lines were obtained but none were homozygous KO (Ξ_2_=4.7, d.f.=1; *P*<0.05). This suggests that ZC3H11A is required for establishing ESC *in vitro*.

### Mice with postnatal *Zc3h11a*-ablation are healthy and viable

We developed an inducible *Zc3h11a*-KO model to assess the effect of *Zc3h11a* ablation postnatally. Loxp-*Zc3h11a* mice were crossed with mice containing fusion of a mutated estrogen receptor T2 and Cre recombinase (Cre-ER), allowing temporal control of floxed gene deletion upon tamoxifen induction *in vivo* [42]. We generated a strain that is homozygous *Zc3h11a*-loxp (*Zc3*^*loxP/loxP*^) with one copy of Cre-ER (CRE.ER^+^ Zc3^loxP/loxP^) and crossed it with the original strain (*Zc3*^*loxP/loxP*^) lacking Cre-ER. The offspring were injected with tamoxifen at week 3-4 after birth (Figure 8A). Genotyping of the tamoxifen-injected mice at week 6 using genomic DNA from tail biopsies revealed a balanced ratio between WT and induced KO (iKO) due to the presence/absence of Cre-ER (Figure 8B). By injecting CRE.ER^+^ Zc3^loxP/loxP^ mice with tamoxifen postnatally we succeeded in achieving >90% reduction of *Zc3h11a* expression in multiple adult tissues including bone marrow, liver and spleen (Figure 8C and D). The examination of tamoxifen-injected mice were carried out at week 12 and involved histology staining of multiple organs including stomach, pancreas, small and large intestine tissues. The histology phenotyping did not exhibit obvious defects between the floxed (WT) and iKO adult mice (Figure 8E and S5). Furthermore, the measurement of body weight, dissected kidney and spleen tissues from WT and inducible ZC3-KO adult mice did not show significant differences (Figure 8F).

**Figure 8.**
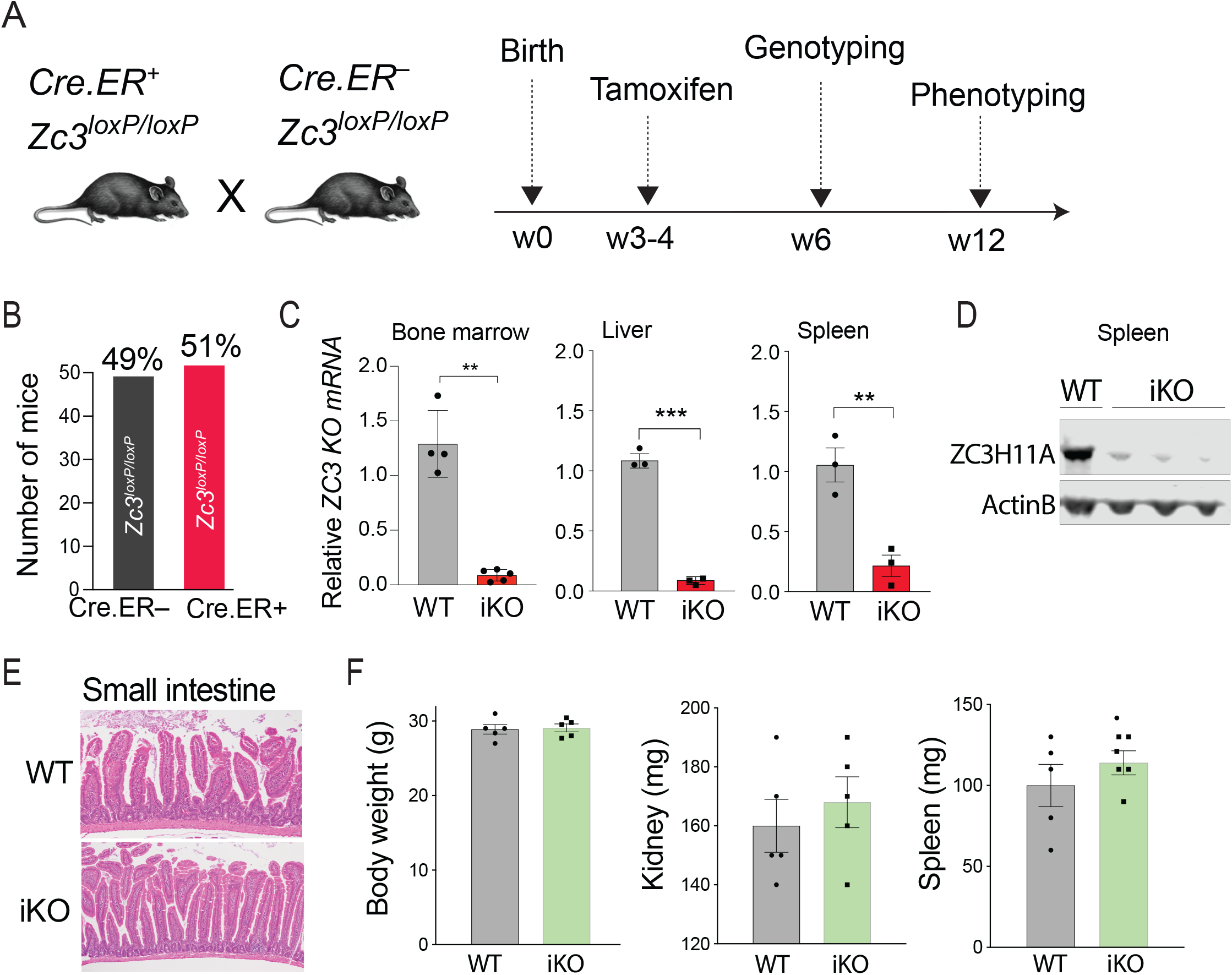
Phenotype characterization of conditional *Zc3h11a*-KO mice. (A) The loxP-*Zc3h11a* mouse model was crossed with mice containing a fusion of a mutated estrogen receptor and Cre recombinase (Cre-ER). The mice were bred to obtain two genotypes of homozygous loxP-*Zc3h11a* mice (Zc3^loxP/loxP^), one with one copy of Cre-ER (CRE.ER^+^ Zc3^loxP/loxP^) and the other with null Cre-ER (CRE.ER^−^ Zc3^loxP/loxP^). These mice were crossed and the offspring were injected with tamoxifen at week 3-4 after birth. The time line indicates the time points of injection and sample collection for genotyping and phenotyping. (B) Genotyping of the Cre-ER Zc3^loxP^ mice. (C) qPCR analysis of *Zc3h11a* mRNA expression in bone marrow, liver and spleen tissues from WT and induced Zc3-KO (iKO) mice both injected with tamoxifen. ** and *** correspond to *t-test P* <0.01 and 0.001, respectively. (D) Western blot analysis of spleen tissues dissected from WT and iKO adult mice. (E) Histology (H&E staining) of small intestine from WT and induced iKO adult mice. (F) Body weight in grams (g), weight of dissected kidney and spleen in milligrams (mg) from WT and induced iKO adult mice. Results are means ± SEM.

## Discussion

ZC3H11A is important for the growth of nuclear replicating viruses, where viruses take advantage of the ZC3H11A protein to facilitate the export of their mRNA transcripts into cytoplasm. Thereby, ZC3H11A is considered a possible target for development of anti-viral therapy. Hence, we developed ZC3H11A mouse models to study its physiological functions across developmental stages. The current study reports that ZC3H11A is an essential protein required for the viability of mouse embryos. Loss of function of ZC3H11A leads to developmental defects and embryo degeneration at peri-implantation stages associated with dysregulation of metabolic pathways such as glycolysis and fatty acid metabolic processes. Interestingly, the defects mainly originate from the epiblast, as most of the down-regulated genes are expressed predominantly in this lineage. Moreover, even though ZC3H11A is expressed in all cells of the blastocyst, *Tdgf1*, one of its key down-regulated target genes is expressed specifically in the epiblast cells [31]. TDGF1 (also called *Cripto*) is a membrane-bound protein, co-receptor for NODAL/GDF3 [43]. TDGF1 and NODAL signaling play important roles during specification of the early lineages and maintenance of the pluripotent epiblast at early post-implantation stages [43]. Interestingly it also controls the metabolic switch occurring at the time of implantation in the mouse, when cells transit from a OXPHOS based metabolism to a glycolytic one [29, 31, 44]. Our CLIP-seq analysis detected two strong peaks for ZC3H11A binding at the 3’ end of the *Tdgf1* mRNA in mESCs (Figure 7E). Furthermore, ZC3H11A binds the 3’ end of *Nup85* and *Pa2g4* mRNA transcripts (Figure 7E). Both *Nup85* and *Pa2g4* were down-regulated in the KO embryos and play crucial roles in embryonic development [38–41]. For instance, NUP85 is a core component of the nuclear pore complex (NPC) proteins and is required for mRNA export and maintenance and assembly of the NPC [40, 41, 45]. Loss of function studies showed that inactivation of the NPC proteins in mouse models resulted in early embryonic lethality [46–48]. Recent phenotypic characterization of the *Nup85* knock-out mouse model from the International Mouse Phenotyping Consortium (www.mousephenotype.org, accessed 22 August 2022) [49] has indicated complete preweaning lethality of *Nup85*^*–/–*^ mice. Furthermore, the ErbB3 binding protein-1 gene (*Ebp1*/*Pa2g4*) is implicated in regulating the proliferation and differentiation during developmental stages. The *Pa2g4* knock-out mice exhibited growth retardation and were 30% smaller than wild-type mice [50]. A recent study has reported more severe phenotypes in *Pa2g4*-deficient mice with death between E13.5 and 15.5, massive apoptosis, and cessation of cell proliferation [38]. These putative ZC3H11A targets identified by CLIP-seq are known to be critical for embryonic viability and implicated in diverse cellular functions, disruption of their expression leads to embryonic degeneration.

Another key down-regulated gene in KO embryos is *Ldha*, the enzyme that controls the level of anaerobic glycolysis by catalyzing the transformation of pyruvate into lactate. Hence, in KO embryos, the establishment of a more anaerobic glycolysis is impaired, which compromises survival when the environment becomes more hypoxic as embryos implant. Upregulation of autophagy as observed in KO embryos can be viewed as reaction to a suboptimal metabolic environment [33]. Although KO embryos can survive up to E6.5, they have already undergone a process of degeneration, as suggested by the upregulation of P53 mediated apoptotic pathway already at E4.5. The transcriptomic analysis also indicated a significant dysregulation in the EMT process (Figure 4C). The EMT process is fundamental for embryo development and takes place during implantation of the embryo into the uterus and during early gastrulation, where embryo is transformed from a single layer to three germ layers. Defects in EMT and subsequently in gastrulation usually lead to a failure in embryonic development [51, 52].

The ZC3H11A protein exhibited strong interactions with members of the RNA-export machinery in ESCs and the top interacting partners with ZC3H11A are members of the TREX complex, including THO proteins (Figure 6). The enrichment analysis of interacting partners with ZC3H11A showed significant enrichment of proteins involved in metabolism of RNA, mRNA 3’-end processing and transport of mature transcript to cytoplasm (Figure S3A). These proteomics results are in agreement with the analysis of the CLIP-seq of ZC3H1A in mESCs that revealed preferential binding at 3’UTRs over the 5’UTRs of target transcripts (Figure 7A). It also supports the model of action that ZC3H11A interacts with TREX-complex proteins and contribute to efficient mRNA maturation and export of the target transcripts. In agreement with this model, several studies have described the pivotal roles of the TREX-complex in the embryonic development [9–11]. THO proteins such as THOC1, THOC2 and THOC5 play essential roles during early development but in a different way than ZC3H11A, as their depletion affects pluripotency establishment and maintenance [9, 10]. In contrast, ZC3H11A depletion does not directly affect pluripotency maintenance. The fact that *Zc3h11a*^*–/–*^ blastocysts did not give rise to ESC lines in the present study may be due to the metabolic impairment rather than a defect in pluripotency maintenance, as they all form outgrowth, in contrast to *Thoc1*^*–/–*^ embryos [10].

Our results provide evidence that ZC3H11A is required for the post-transcriptional regulation of genes that are crucial for the embryonic cell. In contrast to the severe phenotypes in *Zc3h11a* germline KO embryos, *Zc3h11a* inactivation in the adult tissues did not cause obvious defects. The phenotypic characterization of the inducible ZC3-KO adult mice indicated a dispensable role for ZC3H11A in adult tissues and a single surviving *Zc3h11a*^*-/-*^ female showed no pathological conditions, were fertile and gave birth to 10 progeny from three litters. Furthermore, complete inactivation of *Zc3h11a* in human and mouse cell lines did not lead to significant effects on cell growth or viability [1, 3].

## Methods

### Animal models

All mice were group-housed with free access to food and water in the pathogen-free facilities of Uppsala University and INRAE. All procedures described in this study were approved by the Uppsala Ethical Committee on Animal Research (#17346/2017), following the rules and regulations of the Swedish Animal Welfare Agency, and were in compliance with the European Communities Council Directive of 22 September 2010 (2010/63/EU). All efforts were made to minimize animal suffering and to reduce the number of animals used. The loxP *Zc3h11a* mouse model was generated by homologous recombination in mouse C57BL/6 ES cells (Cyagen, USA). The PGK-Cre mice expressing the Cre recombinase in the germ line [53] was obtained as gift from Klas Kullander’s lab (Uppsala University). The CRISPR/cas9 *Zc3h11a* mouse model was purchased from the Mutant Mouse Resource & Research Centers (MMRRC, USA, Strain No: 043457-UCD). For inducible knock-out model, the mice containing fusion of a mutated estrogen receptor T2 and Cre recombinase (Cre-ER) was ordered from The Jackson Laboratory (USA, Strain No: 008463). Mice were genotyped (Tables S2) based on tail biopsies.

### Collection of mouse embryos

The *Zc3h11a* heterozygous males and females were mated and the following day, the presence of a vaginal plug was recorded. To determine the time of developmental lethality, females were sacrificed at E6.5 and embryos dissected out from the decidual swellings. Their morphology was recorded and each of them was then processed for genotyping. Samples for RNA-sequencing were collected at peri-implantation stage (E4). These embryos were bisected using glass needles and both parts were individually snap-frozen. The abembryonic part (mural TE) was used for genotyping and the embryonic part (ICM and polar TE) for subsequent RNA extraction.

### RNA sequencing

The collected embryonic ICM and polar TE (as described above) were used for RNA-seq library preparation using the SMART-Seq HT Kit (Takara Bio USA, Inc.) following the manufacturer’s instructions. Briefly, cDNA was generated using the oligo-dT primer to enrich for mRNA, followed by the tagmentation of the cDNA (Illumina Nextera XT) to generate Illumina-compatible RNA-seq libraries. The libraries were amplified by 12 PCR cycles and size-selected for an average insert size of 150 bp and sequenced as 100 bp paired-end reads using Illumina Nova-Seq. Sequence reads were mapped to the reference mouse genome (mm10) using STAR 2.5.1b [54] with parameter --quantMode GeneCounts to generate read counts. The edgeR (Bioconductor package) [55] was used to identify differentially expressed (DE) genes using gene models for mm10 downloaded from UCSC (www.genome.ucsc.edu). The abundance of gene expression was calculated as count-per-million (CPM) reads. Genes with less than one CPM in at least three samples were filtered out. The filtered libraries were normalized using the trimmed mean of M-values (TMM) normalization method [56]. *P*-values were corrected for multiple testing using the False Discovery Rate (FDR) approach. Gene set enrichment analyses (GSEA) were performed using the fgsea R package [57]. Genes were ranked based on the fold-change and the datasets were downloaded from the GSEA website (https://www.gsea-msigdb.org/gsea/).

### Immunoprecipitation

Mouse embryonic stem cell line (mESC) was cultured on gelatin-coated plates and maintained in Dulbecco’s Modified Eagle Medium (DMEM) complemented with 10% heat-inactivated fetal bovine serum, penicillin (0.2 U/mL), streptomycin (0.2 µg/mL) and L-glutamine (0.2 µg/mL) (Gibco, Waltham, Massachusetts, United States) and supplemented with recombinant mouse Leukemia Inhibitory Factor (LIF, 20 U/ml, Millipore). Cultured mESCs at 60% confluency were washed with PBS twice before the preparation of total lysate. Total protein lysates were prepared using Pierce IP lysis buffer (Thermo Fisher Scientific) supplemented with protease inhibitors (Complete Ultra Tablets, Roche) and Pierce Universal Nuclease (Thermo Fisher Scientific). Lysate was cleared by centrifugation at 20 x *g* for 10 min at 4°C, and incubated rotating end-over-end at 4°C with anti-IgG, anti-ZC3H11A or anti-THOC2 antibodies in Protein LoBind 2-ml tubes (Eppendorf). Thereafter, 30 µg of Dynabeads Protein G (Thermo Fisher Scientific) was added to each tube and incubated for 30 min at room temperature, followed by washing three times with Pierce IP lysis buffer. The co-IP proteins were eluted from the magnetic beads by adding 50 µg of elution buffer (5% SDS, 50mM TEAB, pH 7.55) and heat denaturation for 5 min at 90 °C. The eluted co-IP proteins were used for western blot and Mass spectrometric analysis. co-IP experiments were performed in four replicates.

### Immunoblot analysis

Equal volumes (5 µg) of the prepared co-IPs were separated by SDS-PAGE (4–15%, Bio-Rad) and transferred to PVDF membranes (Millipore). StartingBlock buffer (Thermo Fisher Scientific) was used to block the membrane before the primary anti-ZC3H11A, anti-THOC2 or anti-ALYREF antibodies (1:1000) were added. Proteins were visualized and detected by the Odyssey system (LI-COR).

### Protein clean-up and digestion

The co-IPs were cleaned up and prepared for mass spectrometry quantification using the S-Trap column method [58]. First, the eluted co-IPs were treated by TCEP (5 mM) to reduce disulfide bonds, followed by adding methyl methanethiosulfonate (MMTS) to a final concentration of 15 mM to alkylate cysteines. Thereafter, the lysate was acidified by adding phosphoric acid to a final concentration of 1.2%. The acidified lysate was added to an S-Trap microcolumn (Protifi, Huntington, NY) containing 300 µl of S-Trap buffer (90% MeOH, 100 mM TEAB, pH 7.5) and centrifuged at 4000 × *g* for 2 min. The S-Trap microcolumn was washed twice with S-Trap buffer. The columns were transferred to new tubes and incubated with 10 ng/μL sequencing-grade trypsin (Promega) overnight at 37°C. The digested proteins were eluted by centrifugation at 4000 × *g* for 1 min with 50 mM TEAB, 0.2% formic acid (FA), followed by 50% acetonitrile (ACN)/0.2% FA, and finally 80% ACN/0.1% FA. The eluted peptides were dried down in a vacuum centrifuge (ThermoSavant SPD SpeedVac, Thermo Fisher Scientific), and finally dissolved in 1% FA. Digested peptides were thereafter desalted by StageTips (Thermo Fisher Scientific) according to the manufacturer’s instructions, and subsequently dissolved in 0.1% FA.

### Liquid chromatography and mass spectrometry

The dissolved peptides were quantified by mass spectrometry as previously described [17]. Briefly, a Thermo Scientific EASY-nLC 1000 liquid chromatography system coupled with an Acclaim PepMap 100 (2 cm x 75 μm, 3 μm particles, Thermo Fisher Scientific) pre-column in line with an EASY-Spray PepMap RSLC C18 reversed phase column (50 cm x 75 μm, 2 μm particles, Thermo Fisher Scientific) was utilized to fractionate the peptide samples. The eluted peptides were analyzed on a Thermo Scientific Orbitrap Fusion Tribrid mass spectrometer, operated at a Top Speed data-dependent acquisition scan mode, ion-transfer tube temperature of 275°C, and a spray voltage of 2.0 kV.

### Mass spectrometric data analysis

Data analysis of raw files was performed using MaxQuant software (version 1.6.4) and the Andromeda search engine [59, 60], with the following parameters: cysteine methyl methanethiosulfonate (MMTS) as a static modification and methionine oxidation and protein N-terminal acetylation as variable modifications. First search peptide MS1 Orbitrap tolerance was set to 20 ppm, and iontrap MS/MS tolerance was set to 0.5 Da. Peak lists were searched against the UniProtKB/Swiss-Prot *Mus musculus* proteome database (UP000000589, version 2019-04-01) with a maximum of two trypsin miscleavages per peptide. The MaxQuant contaminants database was also utilized. A decoy search was made against the reversed database, with the peptide and protein false discovery rates both set to 1%. Only proteins identified with at least two peptides of at least 7 amino acids in length were considered reliable. The peptide output from MaxQuant was filtered by removing reverse database hits, potential contaminants, and proteins only identified by site (PTMs).

Intensity values were used to determine the protein abundance. First, proteins with missing values in more than one replicate in at least one group were filtered out. Thereafter, the filtered intensities were normalized using the variance stabilizing normalization (vsn) method [61] and followed by the imputation of missing values with the deterministic minimal value approach (MinDet) [62] to replace the missing values in the normalized intensities. The normalized intensities were fitted to a linear model and the empirical Bayes moderated t-statistics and their associated *P*-values were used to calculate the significance of differential enriched proteins [63, 64]. The *P*-values were adjusted for multiple testing using the Benjamini–Hochberg procedure [65]. Proteomics data was visualized using the ggplot R-package and Cytoscape v3.8.2.

### Crosslinking immunoprecipitation sequencing (CLIP-seq)

Cultured mESCs were cross-linked using a 254 nM UV crosslinker with an energy setting of 400 mJ/cm^2^. The cross-linked cells were collected in ice-cold PBS with a cell scraper and aliquoted in 1.5 ml tubes (25 million cells/ml). The CLIP-seq library preparation was performed as indicated [66]. Briefly, the total lysate was digested with RNase-I and immunoprecipitated with 10 µg of the following antibodies: anti-ZC3H11A (HPA028526 and HPA028490, Atlas Antibodies), anti-ALYREF (ab202894, Abcam) or anti-IgG (ab37415, Abcam). The IP-RNA complexes were loaded on a 4–12% Bis-Tris (Bio-Rad) gel, transferred to a nitrocellulose membrane, and the bands above 75 kDa for each lane were cut. Extracted RNA molecules from the membrane were used for Illumina library construction as indicated [66]. CLIP-seq reads were trimmed out using *trim_galore* with the criteria to remove reads with low quality and shorter than 15 bp. The trimmed reads were aligned to the mouse reference genome mm10 using STAR aligner with end-to-end options --alignEndsType EndToEnd. The CLAM workflow were used for peak calling and counting the fold enrichment of IP vs IgG control [67]. The identified peaks with adjusted *P*-value <0.01 were annotated to the mouse mm10 genome using the *peak_annotator* function from CLAM. HOMER software was used for motif finding using the findMotifsGenome.pl script with default parameters for RNA motifs [68].

### Quantitative RT-PCR

Total RNA was extracted using the RNeasy Mini kit (Qiagen) and the samples were treated with DNase I to eliminate genomic DNA. The High Capacity cDNA Reverse Transcription Kit (Applied Biosystems) was used to generate cDNA from RNA. Quantitative PCR analysis was performed in ABI MicroAmp Optical 384-well Reaction plates on an ABI 7900 real-time PCR instrument using SYBR gene expression reagents (Applied Biosystems). The amplification and detection of each gene was performed using forward and reverse primers for *Zc3h11a*, F: TGCCTAATCAGGGAGAAGACTG, R: AGCTTCACAGTGACGGAATG and *Actb* as a housekeeping gene F: CTAAGGCCAACCGTGAAAAG, R: ATCACAATGCCTGTGGTACG.

### Derivation of ESCs

E3.5 blastocysts were collected from heterozygous matings. They were plated individually on a layer of Mitomycine C inactivated mouse embryonic fibroblasts (feeder layer) in 4-well plates, in naïve ESC medium. This medium was composed of Chemically Defined Medium (CDM) supplemented with LIF (700 U/ml), PD0332552 (1 µM) and CHIR99201 (final 3 µM) (2i/Lif CDM) [69]. After 4-5 days, the blastocysts have attached and outgrowths formed. Individual outgrowths were dislodged from the feeders, dissociated in Tryple Select (Invitrogen) and the single cell suspension was plated in 4-well plates on fresh feeders. ESC colonies appeared within the following days and were individually picked, dissociated and replated on feeders in 2i/Lif CDM. The procedure was repeated a few times, until stable expansion of the ESCs that allows passaging using trypsin and removal of feeders, replaced by plate coating with gelatin 0.2% and serum.

### Immunostaining of ZC3H11A on pre-implantation embryos

Mouse CD1 embryos were collected at 4-cell (E1.5) and blastocyst (E3.5) stage in M2 medium (Sigma) by oviduct and uterine flushing, respectively. They were fixed in 2% PFA for 20 min, followed by permeabilization by 1% Triton-×100, for 30 min. Permeabilized samples were blocked with 1% BSA in PBS for 40 min, followed by incubation with primary antibodies in 1% BSA overnight at 4°C. The day after, samples were washed by PBS and incubated with the fluorophore conjugated secondary antibodies (Jackson ImmunoResearch) for an hour at room temperature. Samples were then washed and stained by DAPI for nuclei staining. Samples were mounted in Vectashield mounting agent (Vectorlabs, H1000). Embryos were imaged by a Zeiss LSM710 confocal microscope. Antibodies used were anti-SC35 (ab11826, Abcam, 1:250) and anti-ZC3H11A (HPA028526, Atlas Antibodies, 1:300)

## Author contributions

SY and LA conceived the study. SY performed experimental and bioinformatic analysis. AJ, VB and JO performed experiments on embryos and ESC derivation with contribution from SY. SY and ML performed mass spectrometry analysis. SY, AJ and LA wrote the paper with input from ML. All authors approved the final version before submission.

## Acknowledgments

This project was funded by the Swedish Research Council (2017-02907) and the Knut and Alice Wallenberg Foundation (KAW 2017.0071), as well as by the French REVIVE Labex (Investissement d’Avenir, ANR-10-LABX-73). Sequencing was performed by the SNP&SEQ Technology Platform in Uppsala. The Swedish facility is part of the National Genomics Infrastructure (NGI) Sweden and Science for Life Laboratory. The SNP&SEQ Platform is also supported by the Swedish Research Council and the Knut and Alice Wallenberg Foundation. Animal experiments performed in France were done in INRAE Infectiology of Fishes and Rodents Facility (IERP-UE907, Jouy-en-Josas Research Center), which belongs to the National Distributed Research Infrastructure for the Control of Animal and Zoonotic Emerging Infectious Diseases through In Vivo Investigation (EMERG’IN, doi: 10.15454/1.5572352821559333E12). Fluorescent images were acquired in the French ISC MIMA2 (Microscopy and Imaging Facility for Microbes, Animals and Foods, doi: 10.15454/1.5572348210007727E12).

## Additional information

### Ethics statement

Animal procedures were carried out according to the rules and regulations of the Swedish Animal Welfare Agency and French national rules on Ethics and Animal Welfare in the Animal Facility; and were in compliance with the European Communities Council Directive of 22 September 2010 (2010/63/EU). This work was approved by the French Ministry of Higher Education, Research, and Innovation (n°15-55 &21-01) and the local Ethical Committee (INRAE Jouy-en-Josas Centre). The study was carried out in compliance with the ARRIVE guidelines.

### Data availability

The mass spectrometry proteomics data have been deposited to the ProteomeXchange Consortium via the PRIDE partner repository with the accession numbers (to be added). The RNA-seq reads have been submitted to the sequence read archive (http://www.ncbi.nlm.nih.gov/sra) with the accession numbers (to be added).

## Supplementary information

**Figure S1.**
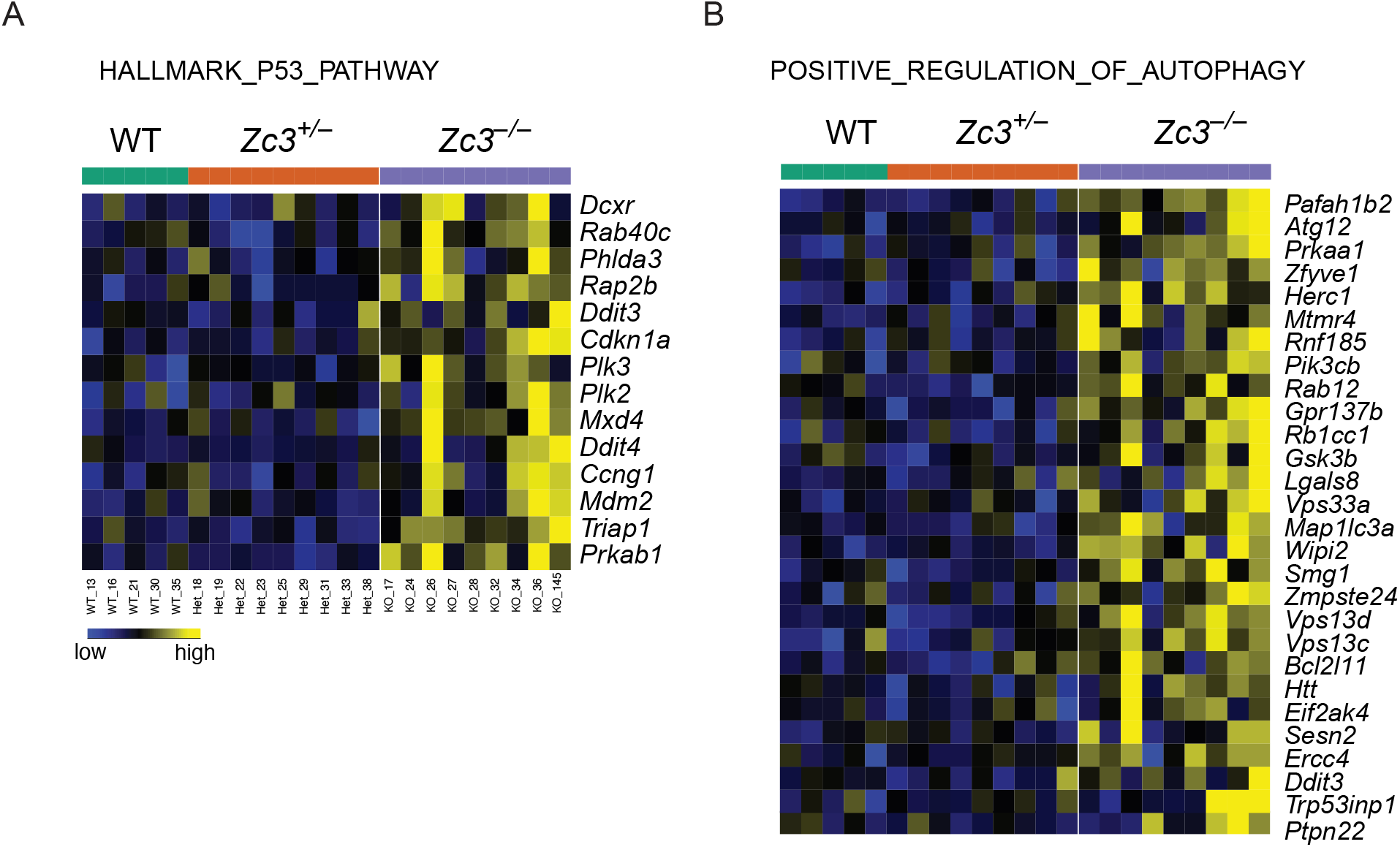
Heatmaps of the normalized expression of genes involved in the P53 pathway (A) and in positive regulation of autophagy (B).

**Figure S2.**
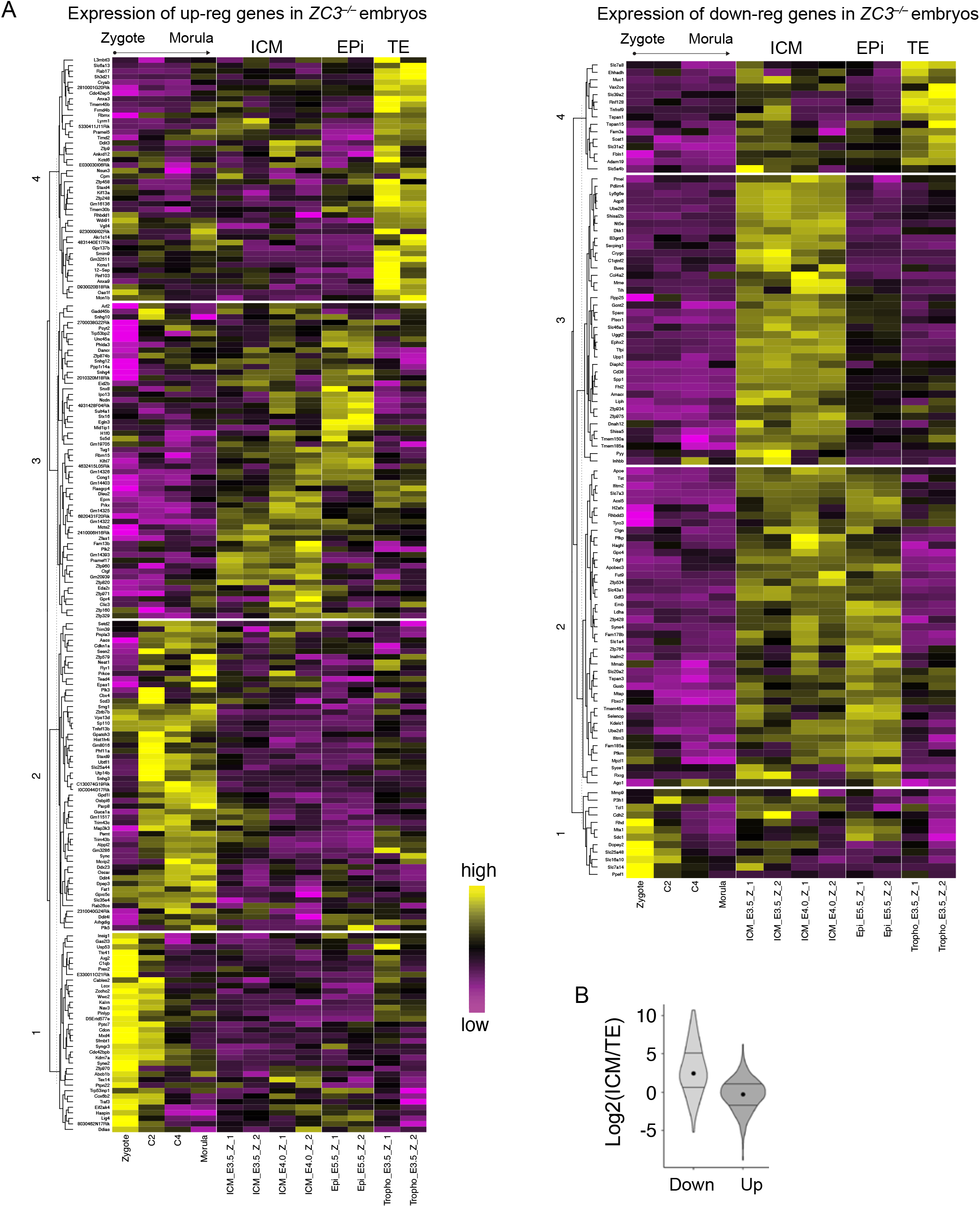
(A) Heatmap of up-regulated genes (A, left) and down-regulated genes (A, right) in *Zc3h11a*^−/–^ embryos (FDR <0.05) and their expression profile during embryonic stages as indicated. Re-analyzed data from GSE76505 and E-MTAB-2950. (B) Violin plot of the relative expression of inner cell mass (ICM)-related genes and trophectoderm (TE)-related genes (ICM/TE) as detected among the down-regulated and up-regulated genes in *Zc3h11a*^−/–^ embryos (FDR <0.05).

**Figure S3.**
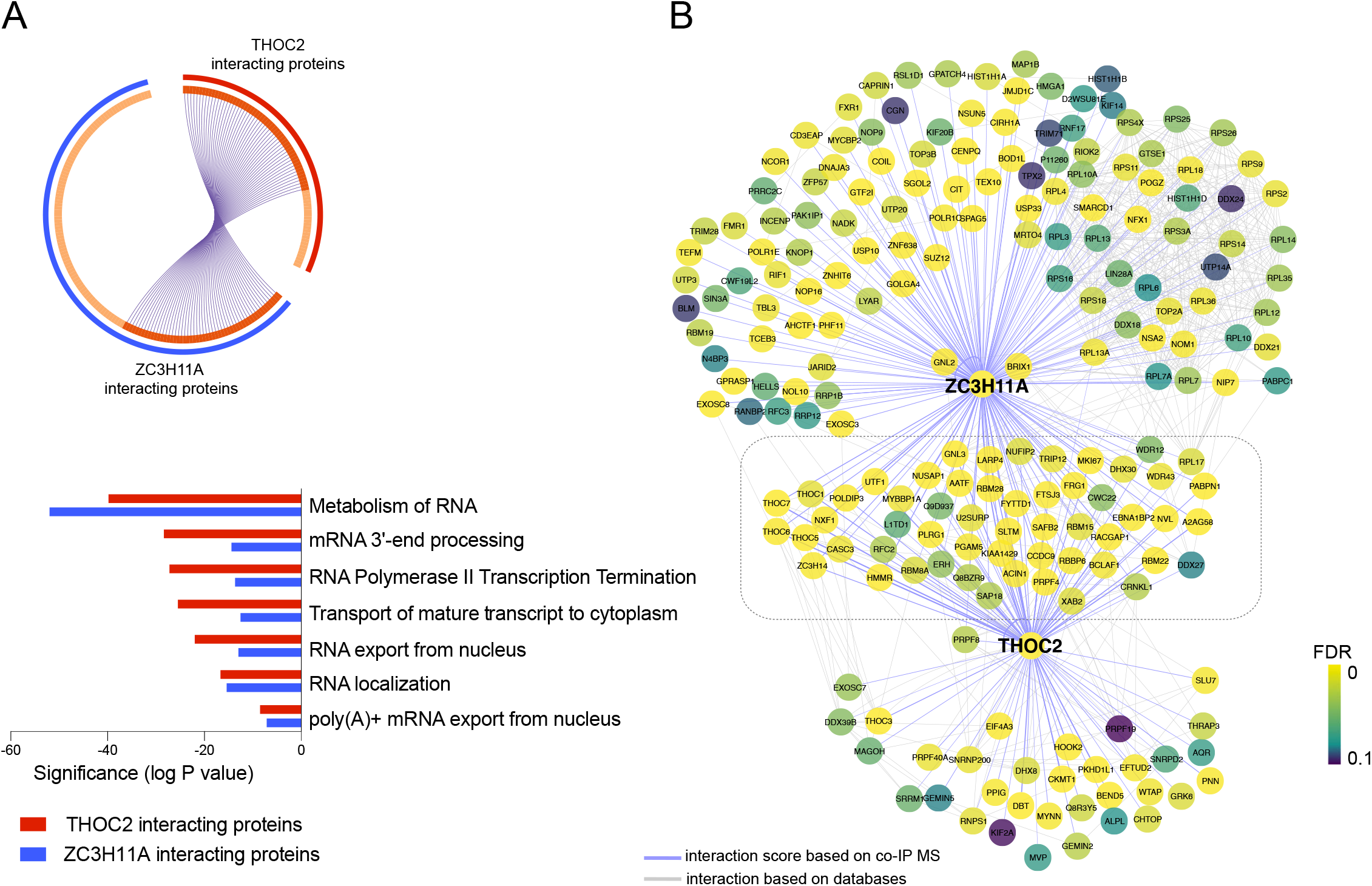
(A, top) Chord diagram illustrating the overlap between the interacting partners detected with ZC3H11A and THOC2 co-IPs. (A, bottom) GO analysis of the identified interacting partners with ZC3H11A and THOC2 co-IPs in mESCs. (B) Network analysis of the identified interacting partners with ZC3H11A and THOC2 based on co-IPs in mESCs. Blue lines represent the interaction detected by our proteomics analysis, and gray lines represent the predicted interaction from the STRING database.

**Figure S4.**
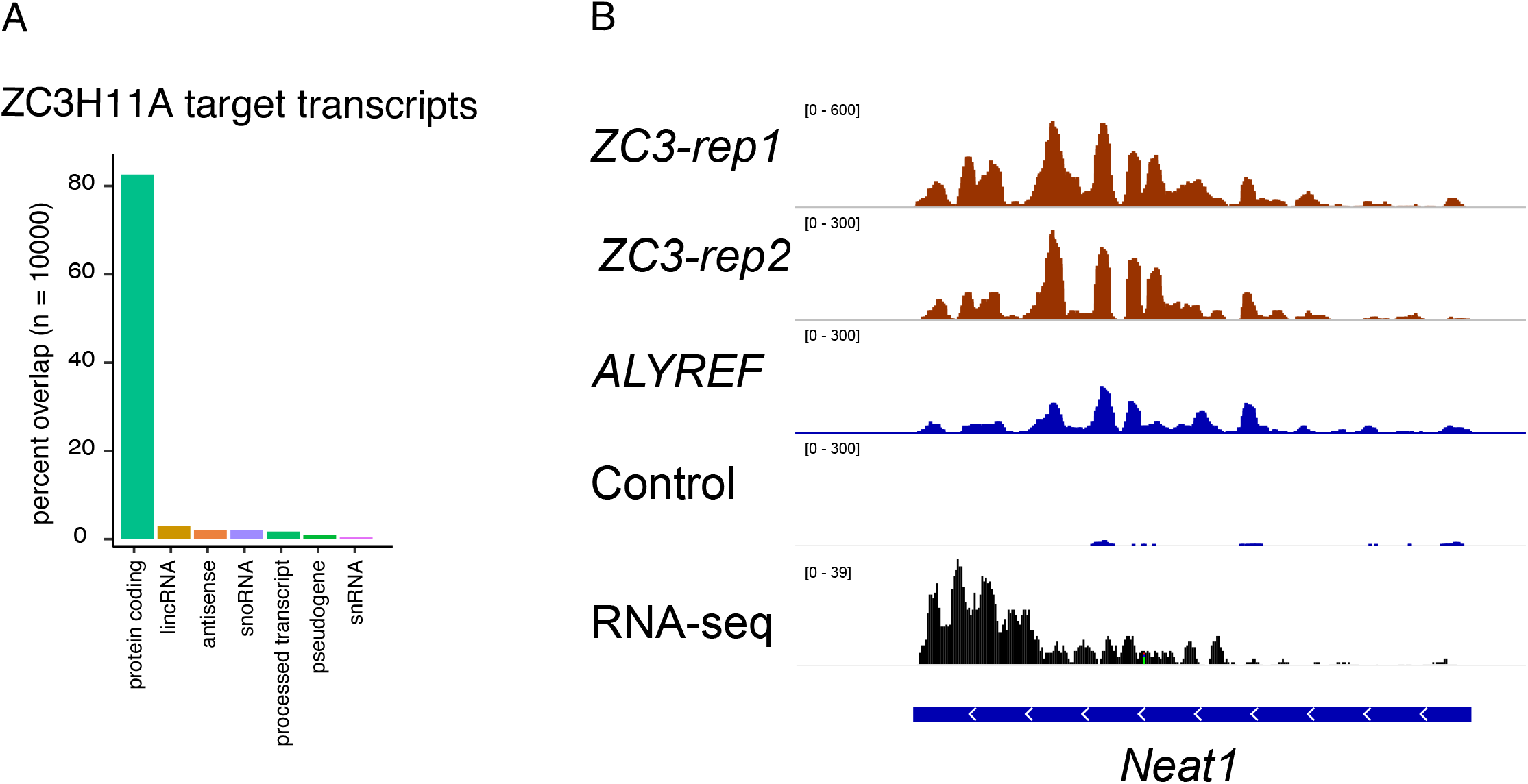
(A) Genome-wide distribution of the ZC3H11A CLIP-seq peaks from mESCs across different types of transcripts. (B) Visualization of CLIP-seq reads and their distribution over the *Neat1* gene.

**Figure S5.**
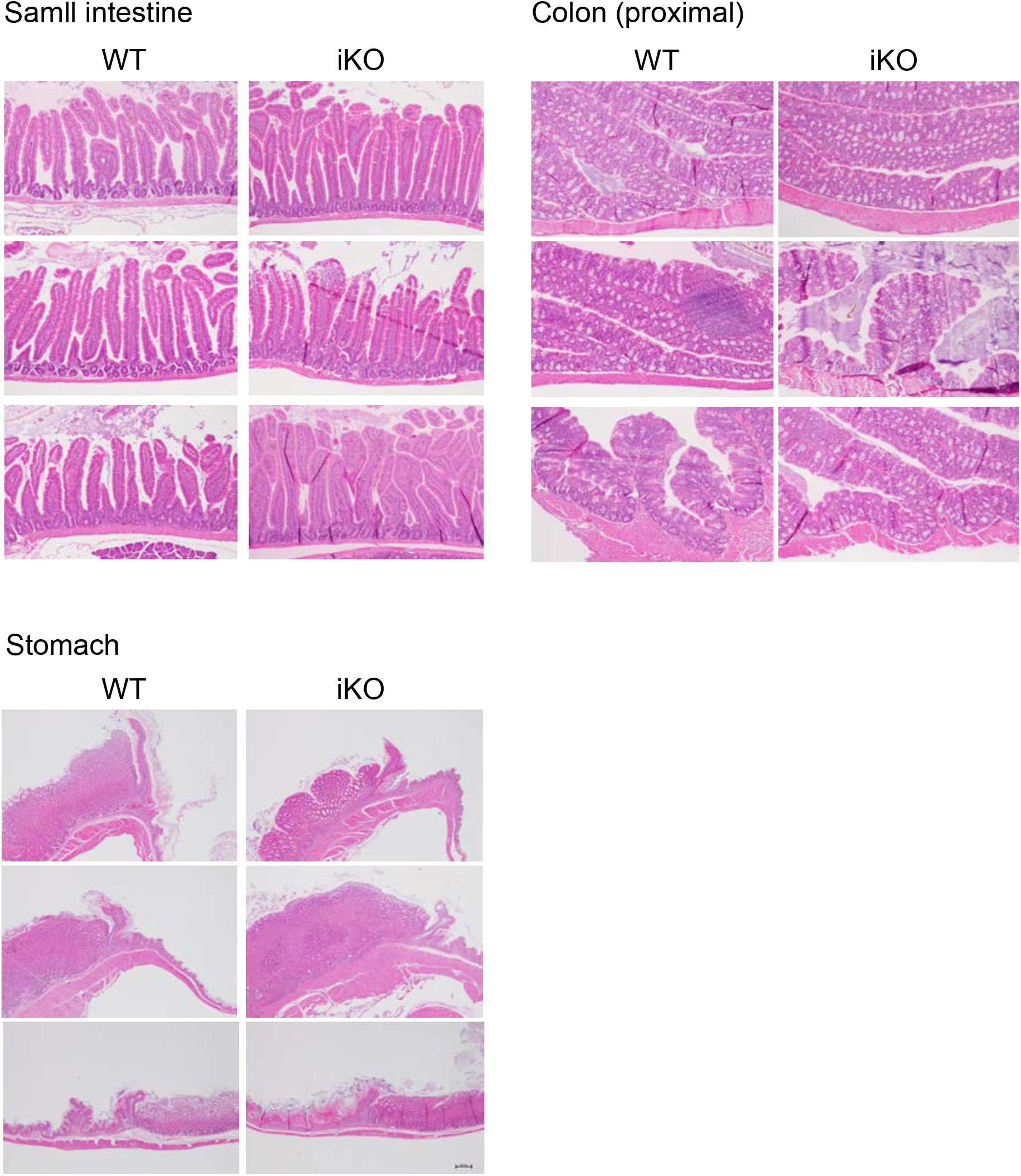
Histology staining of intestine, colon and stomach tissues from three WT and three induced ZC3-KO adult mice.

**Table S1**. Differentially expressed genes in *Zc3h11a* knock-out (KO) E4.5 embryos.

**Table S2**. Sequences of PCR primers used for mouse genotyping.

## References

1. S. Younis, et al., Multiple nuclear-replicating viruses require the stress-induced protein ZC3H11A for efficient growth. Proc. Natl. Acad. Sci. U. S. A. 115, E3808–E3816 (2018).

2. E. G. Folco, C.-S. Lee, K. Dufu, T. Yamazaki, R. Reed, The proteins PDIP3 and ZC11A associate with the human TREX complex in an ATP-dependent manner and function in mRNA export. PLoS One 7, e43804 (2012).

3. L. Yang, et al., Porcine ZC3H11A Is Essential for the Proliferation of Pseudorabies Virus and Porcine Circovirus 2. ACS Infect. Dis. (2022) https://doi.org/10.1021/ACSINFECDIS.2C00150.

4. C. G. Heath, N. Viphakone, S. A. Wilson, The role of TREX in gene expression and disease. Biochem. J. 473, 2911–2935 (2016).

5. B. Chi, et al., Aly and THO are required for assembly of the human TREX complex and association of TREX components with the spliced mRNA. Nucleic Acids Res. 41, 1294–1306 (2013).

6. K. Dufu, et al., ATP is required for interactions between UAP56 and two conserved mRNA export proteins, Aly and CIP29, to assemble the TREX complex. Genes Dev. 24, 2043–53 (2010).

7. M. Y. Hein, et al., A Human Interactome in Three Quantitative Dimensions Organized by Stoichiometries and Abundances. Cell 163, 712–723 (2015).

8. L. Ding, et al., A Genome-Scale RNAi Screen for Oct4 Modulators Defines a Role of the Paf1 Complex for Embryonic Stem Cell Identity. Cell Stem Cell 4, 403–415 (2009).

9. L. Wang, et al., The THO Complex Regulates Pluripotency Gene mRNA Export and Controls Embryonic Stem Cell Self-Renewal and Somatic Cell Reprogramming. Cell Stem Cell 13, 676–690 (2013).

10. X. Wang, Y. Chang, Y. Li, X. Zhang, D. W. Goodrich, Thoc1/Hpr1/p84 Is Essential for Early Embryonic Development in the Mouse. Mol. Cell. Biol. 26, 4362–4367 (2006).

11. A. Mancini, et al., THOC5/FMIP, an mRNA export TREX complex protein, is essential for hematopoietic primitive cell survival in vivo. BMC Biol. 8, 1–17 (2010).

12. V. O. Wickramasinghe, R. A. Laskey, Control of mammalian gene expression by selective mRNA export. Nat. Rev. Mol. Cell Biol. 16, 431–42 (2015).

13. S. Younis, et al., The ZBED6-IGF2 axis has a major effect on growth of skeletal muscle and internal organs in placental mammals. Proc. Natl. Acad. Sci. U. S. A. 115, E2048–E2057 (2018).

14. X. Wang, et al., ZBED6 negatively regulates insulin production, neuronal differentiation, and cell aggregation in MIN6 cells. FASEB J. 33, 88–100 (2019).

15. R. Naboulsi, M. Larsson, L. Andersson, S. Younis, ZBED6 regulates Igf2 expression partially through its regulation of miR483 expression. Sci. Rep. 11 (2021).

16. X. Wang, et al., ZBED6 counteracts high-fat diet-induced glucose intolerance by maintaining beta cell area and reducing excess mitochondrial activation. Diabetologia 64, 2292–2305 (2021).

17. S. Younis, et al., The importance of the ZBED6-IGF2 axis for metabolic regulation in mouse myoblast cells. FASEB J. 34, 10250–10266 (2020).

18. M. A. Ali, et al., Transcriptional modulator ZBED6 affects cell cycle and growth of human colorectal cancer cells. Proc. Natl. Acad. Sci. U. S. A. 112, 7743–7748 (2015).

19. Q. Deng, D. Ramsköld, B. Reinius, R. Sandberg, Single-cell RNA-seq reveals dynamic, random monoallelic gene expression in mammalian cells. Science 343, 193–196 (2014).

20. S. Kanungo, K. Wells, T. Tribett, A. El-Gharbawy, Glycogen metabolism and glycogen storage disorders. Ann. Transl. Med. 6, 474–474 (2018).

21. H. Wakabayashi, M. Tsuchiya, K. Yoshino, K. Kaku, H. Shigei, Hereditary deficiency of lactate dehydrogenase H-subunit. Intern. Med. 35, 550–554 (1996).

22. J. K. Reddy, S. K. Goel, M. R. Nemali, Transcriptional regulation of peroxisomal fatty acyl-CoA oxidase and enoyl-CoA hydratase/3-hydroxyacyl-CoA dehydrogenase in rat liver by peroxisome proliferators. Proc. Natl. Acad. Sci. U. S. A. 83, 1747–1751 (1986).

23. C. Qi, et al., Absence of spontaneous peroxisome proliferation in enoyl-CoA hydratase/L-3-hydroxyacyl-CoA dehydrogenase-deficient mouse liver: Further support for the role of fatty acyl CoA oxidase in PPARα ligand metabolism. J. Biol. Chem. 274, 15775–15780 (1999).

24. O. Lieven, J. Knobloch, U. Rüther, The regulation of Dkk1 expression during embryonic development. Dev. Biol. 340, 256–268 (2010).

25. H. Suzuki, et al., Structural basis of the autophagy-related LC3/Atg13 LIR complex: Recognition and interaction mechanism. Structure 22, 47–58 (2014).

26. Y. Zhang, et al., Dynamic epigenomic landscapes during early lineage specification in mouse embryos. Nat. Genet. 2017 501 50, 96–105 (2017).

27. K. K. Abe, et al., The first murine zygotic transcription is promiscuous and uncoupled from splicing and 3′ processing. EMBO J. 34, 1523–1537 (2015).

28. C. Chen, et al., The Vg1-related protein Gdf3 acts in a Nodal signaling pathway in the pre-gastrulation mouse embryo. Development 133, 319–329 (2006).

29. M. T. Johnson, S. Mahmood, M. S. Patel, Intermediary metabolism and energetics during murine early embryogenesis. J. Biol. Chem. 278, 31457–31460 (2003).

30. A. Malkowska, C. Penfold, S. Bergmann, T. E. Boroviak, A hexa-species transcriptome atlas of mammalian embryogenesis delineates metabolic regulation across three different implantation modes. Nat. Commun. 13, 1–12 (2022).

31. A. Fiorenzano, et al., Cripto is essential to capture mouse epiblast stem cell and human embryonic stem cell pluripotency. Nat. Commun. 7 (2016).

32. H. Yan, et al., Fatty acid oxidation is required for embryonic stem cell survival during metabolic stress. EMBO Rep. 22 (2021).

33. R. C. Russell, H. X. Yuan, K. L. Guan, Autophagy regulation by nutrient signaling. Cell Res. 24, 42–57 (2014).

34. U. Kühn, E. Wahle, Structure and function of poly(A) binding proteins. Biochim. Biophys. Acta - Gene Struct. Expr. 1678, 67–84 (2004).

35. G. M. Hautbergue, et al., UIF, a New mRNA Export Adaptor that Works Together with REF/ALY, Requires FACT for Recruitment to mRNA. Curr. Biol. 19, 1918–1924 (2009).

36. M. Shi, et al., ALYREF mainly binds to the 5′ and the 3′ regions of the mRNA in vivo. Nucleic Acids Res. 45, 9640–9653 (2017).

37. H. Zhang, et al., DEAD-Box Helicase 18 Counteracts PRC2 to Safeguard Ribosomal DNA in Pluripotency Regulation. Cell Rep. 30, 81-97.e7 (2020).

38. H. R. Ko, et al., Roles of ErbB3-binding protein 1 (EBP1) in embryonic development and gene-silencing control. Proc. Natl. Acad. Sci. U. S. A. 116, 24852–24860 (2019).

39. K. M. Neilson, et al., Pa2G4 is a novel Six1 co-factor that is required for neural crest and otic development. Dev. Biol. 421, 171–182 (2017).

40. A. Harel, et al., Removal of a single pore subcomplex results in vertebrate nuclei devoid of nuclear pores. Mol. Cell 11, 853–864 (2003).

41. T. C. Walther, et al., The conserved Nup107-160 complex is critical for nuclear pore complex assembly. Cell 113, 195–206 (2003).

42. A. Ventura, et al., Restoration of p53 function leads to tumour regression in vivo. Nat. 2006 4457128 445, 661–665 (2007).

43. C. Bianco, et al., Role of Cripto-1 in Stem Cell Maintenance and Malignant Progression. Am. J. Pathol. 177, 532–540 (2010).

44. W. Zhou, et al., HIF1α induced switch from bivalent to exclusively glycolytic metabolism during ESC-to-EpiSC/hESC transition. EMBO J. 31, 2103–2116 (2012).

45. A. L. Goldstein, C. A. Snay, C. V. Heath, C. N. Cole, Pleiotropic nuclear defects associated with a conditional allele of the novel nucleoporin Rat9p/Nup85p. Mol. Biol. Cell 7, 917–934 (1996).

46. M. Smitherman, K. Lee, J. Swanger, R. Kapur, B. E. Clurman, Characterization and Targeted Disruption of Murine Nup50, a p27 Kip1 -Interacting Component of the Nuclear Pore Complex. Mol. Cell. Biol. 20, 5631–5642 (2000).

47. K. Okita, et al., Targeted disruption of the mouse ELYS gene results in embryonic death at peri-implantation development. Genes to Cells 9 (2004).

48. J. Van Deursen, J. Boer, L. Kasper, G. Grosveld, G2 arrest and impaired nucleocytoplasmic transport in mouse embryos lacking the proto-oncogene CAN/Nup214. EMBO J. 15, 5574–5583 (1996).

49. M. E. Dickinson, et al., High-throughput discovery of novel developmental phenotypes. Nature 537, 508– 514 (2016).

50. Y. Zhang, et al., Alterations in cell growth and signaling in ErbB3 binding protein-1 (Ebp1) deficient mice. BMC Cell Biol. 9 (2008).

51. H. Acloque, M. S. Adams, K. Fishwick, M. Bronner-Fraser, M. A. Nieto, Epithelial-mesenchymal transitions: the importance of changing cell state in development and disease. J. Clin. Invest. 119, 1438 (2009).

52. T. Chen, Y. You, H. Jiang, Z. Z. Wang, Epithelial–mesenchymal transition (EMT): A biological process in the development, stem cell differentiation, and tumorigenesis. J. Cell. Physiol. 232, 3261–3272 (2017).

53. Y. Lallemand, V. Luria, R. Haffner-Krausz, P. Lonai, Maternally expressed PGK-Cre transgene as a tool for early and uniform activation of the Cre site-specific recombinase. Transgenic Res. 7, 105–112 (1998).

54. A. Dobin, et al., STAR: Ultrafast universal RNA-seq aligner. Bioinformatics 29, 15–21 (2013).

55. M. D. Robinson, D. J. McCarthy, G. K. Smyth, edgeR: A Bioconductor package for differential expression analysis of digital gene expression data. Bioinformatics 26, 139–140 (2009).

56. M. D. Robinson, A. Oshlack, A scaling normalization method for differential expression analysis of RNA-seq data. Genome Biol. 11, R25 (2010).

57. A. A. Sergushichev, An algorithm for fast preranked gene set enrichment analysis using cumulative statistic calculation. bioRxiv, 060012 (2016).

58. K. R. Ludwig, M. M. Schroll, A. B. Hummon, Comparison of In-Solution, FASP, and S-Trap Based Digestion Methods for Bottom-Up Proteomic Studies. J. Proteome Res. 17, 2480–2490 (2018).

59. J. Cox, M. Mann, MaxQuant enables high peptide identification rates, individualized p.p.b.-range mass accuracies and proteome-wide protein quantification. Nat. Biotechnol. 26, 1367–1372 (2008).

60. S. Tyanova, T. Temu, J. Cox, The MaxQuant computational platform for mass spectrometry-based shotgun proteomics. Nat. Protoc. 11, 2301–2319 (2016).

61. W. Huber, A. Von Heydebreck, H. Sültmann, A. Poustka, M. Vingron, Variance stabilization applied to microarray data calibration and to the quantification of differential expression in Bioinformatics, (Oxford University Press, 2002), pp. S96–S104.

62. C. Lazar, L. Gatto, M. Ferro, C. Bruley, T. Burger, Accounting for the Multiple Natures of Missing Values in Label-Free Quantitative Proteomics Data Sets to Compare Imputation Strategies. J. Proteome Res. 15, 1116–1125 (2016).

63. X. Zhang, et al., Proteome-wide identification of ubiquitin interactions using UbIA-MS. Nat. Protoc. 13, 530–550 (2018).

64. G. K. Smyth, Linear Models and Empirical Bayes Methods for Assessing Differential Expression in Microarray Experiments. Stat. Appl. Genet. Mol. Biol. 3, 1–25 (2004).

65. Y. Benjamini, Y. Hochberg, Controlling the False Discovery Rate: A Practical and Powerful Approach to Multiple Testing. J. R. Stat. Soc. Ser. B 57, 289–300 (1995).

66. E. L. Van Nostrand, et al., “Robust, cost-effective profiling of RNA binding protein targets with single-end enhanced crosslinking and immunoprecipitation (SeCLIP)” in Methods in Molecular Biology, (2017), pp. 177–200.

67. Z. Zhang, Y. Xing, CLIP-seq analysis of multi-mapped reads discovers novel functional RNA regulatory sites in the human transcriptome. Nucleic Acids Res. 45, 9260–9271 (2017).

68. S. Heinz, et al., Simple Combinations of Lineage-Determining Transcription Factors Prime cis-Regulatory Elements Required for Macrophage and B Cell Identities. Mol. Cell 38, 576–589 (2010).

69. M. Tosolini, A. Jouneau, From naive to primed pluripotency: In vitro conversion of mouse embryonic stem cells in epiblast stem cells. Methods Mol. Biol. 1341, 209–216 (2015).

